# TORC2 dependent phosphorylation modulates calcium regulation of fission yeast myosin

**DOI:** 10.1101/498865

**Authors:** Karen Baker, Irene A. Gyamfi, Gregory I. Mashanov, Justin E. Molloy, Michael A. Geeves, Daniel P. Mulvihill

## Abstract

All cells have the ability to respond to changes in their environment. Signalling networks modulate cytoskeleton and membrane organisation to impact cell cycle progression, polarised cell growth and multicellular development according to the environmental setting. Using diverse *in vitro, in vivo* and single molecule techniques we have explored the role of myosin-1 signalling in regulating endocytosis during both mitotic and meiotic cell cycles. We have established that a conserved serine within the neck region of the sole fission yeast myosin-1 is phosphorylated in a TORC2 dependent manner to modulate myosin function. Myo1 neck phosphorylation brings about a change in the conformation of the neck region and modifies its interaction with calmodulins, Myo1 dynamics at endocytic foci, and promotes calcium dependent switching between different calmodulin light chains. These data provide insight into a novel mechanism by which myosin neck phosphorylation modulates acto-myosin dynamics to control polarised cell growth in response to mitotic and meiotic cell-cycle progression and the cellular environment.

## Introduction

The actin cytoskeleton underpins cellular organisation by maintaining cell shape and through the transmission of mechanical signals between the cell periphery and nucleus, to influence protein expression, organisation and cellular architecture in response to needs of the cell. Myosins, actin-associated motor-proteins, work in collaboration to facilitate global cytoskeletal organisation and a plethora of transport processes including cell migration, intracellular transport, tension sensing and cell division (O’Connell *et al*, 2007). While there are many classes of myosin, each contains an actin binding ATPase motor domain, which exerts force against actin, a lever arm or neck region that contains light chain binding IQ motifs, and a tail region which specifies cargo binding and other molecular interactions.

Although different classes of myosin perform very different cellular functions they all operate by the same basic mechanism, whereby the motor domain undergoes cyclical interactions with actin coupled to the breakdown of ATP. Each molecule of ATP that is converted to ADP and inorganic phosphate can generate movement along actin of between 5–25 nm and force of up to 5 pN. Regulation of acto-myosin motility is multi-faceted (Heissler & Sellers, 2016a), combining regulatory pathways operating via the actin track (historically called thin-filament regulation), or myosin-linked regulation (historically called thick filament regulation) which is often mediated via phosphorylation of the heavy chain or light chain(s) or by calcium-regulation of light chain binding (Heissler & Sellers, 2016b). It has been shown that phosphorylation at the conserved “TEDS” motif within the myosin motor domain of class 1 myosin affects acto-myosin interaction (Bement & Mooseker, 1995); phosphorylation within the tail region of class 5 myosin controls cargo binding (Rogers *et al*, 1999), whereas phosphorylation of class 2 myosin light chains and/or heavy chain can change the folded state of the heavy chain, affecting both actin interaction and ability to form filaments (Redowicz, 2001; Kendrick-Jones *et al*, 1987; Pasapera *et al*, 2015). So, phosphoregulation of myosin can occur in the head, neck and tail regions and also the light chains and its effects are manifold and vary across myosin classes and between paralogues within the same class. Its effect on motile function is still not fully understood for many myosins, especially within yeast (East & Mulvihill, 2011).

The fission yeast, *Schizosaccharomyces pombe*, genome encodes for 5 myosin heavy chains from classes 1, 2, and 5 (Win *et al*, 2002), representing the basic subset of these actin-associated motor proteins found in eukaryotic cells. The single class 1 myosin, Myo1, is a 135 kDa protein, with motor domain, neck region (with two canonical IQ motifs) and a 49 kDa tail region containing a, so-called, tail-homology-2 domain, PH domain, SH3 domain and a carboxyl-terminal acidic region that associates with and activates the Arp2/3 complex to nucleate actin polymerisation (Lee *et al*, 2000). The myosin motor has a conserved TEDS site, phosphorylated by a Ste20 protein kinase, to modulate the protein’s ability to associate with actin (Attanapola *et al*, 2009). Myo1 associates with membranes, primarily at sites of cell growth, where it is required for endocytosis, actin organisation and spore formation (Sirotkin *et al*, 2005; Lee *et al*, 2000; Itadani *et al*, 2007).

Calmodulin or calmodulin-like light chains associate with the IQ motifs within the myosin neck, providing a mechanism to regulate the length and stiffness of the lever arm (Trybus *et al*, 2007) and behaviour of the motor domain (Adamek *et al*, 2008). Calmodulins are ubiquitous calcium binding proteins that associate with and regulate the cellular function of diverse proteins. Calcium associates with up to four EF hand motifs within the calmodulin molecule to bring about a change in its conformation to modulate its affinity for IQ motifs within binding partner proteins (Crivici & Ikura, 1995). S. *pombe* encodes for two calmodulin like proteins, Cam1 and Cam2 (Takeda & Yamamoto, 1987; Itadani *et al*, 2007). Cam1 is a typical calmodulin that associates with IQ domain containing proteins in a calcium dependent manner, to affect functions as diverse as endocytosis, spore formation, cell division or maintaining spindle pole body integrity (Takeda & Yamamoto, 1987; Moser *et al*, 1995; 1997; Itadani *et al*, 2010). Unlike Cam1, Cam2 is not essential and is predicted to be insensitive to calcium, however like Cam1 it has been reported to regulate Myo1 (Sammons *et al*, 2011; Itadani *et al*, 2007). While cells lacking Cam2 show defects in spore formation they have no significant growth-associated phenotypes during the vegetative growth cycle.

TOR (Target of Rapamycin) signaling plays a key role in modulating cell growth in response to changes in cell cycle status and environmental conditions (Laplante & Sabatini, 2012). The mTOR kinase forms two distinct protein complexes TOR complex 1 (TORC1) and TORC2, each defined by unique components that are highly conserved across species. While both TORC1 and TORC2 have been implicated in the control of cell migration and F-actin organisation (Liu & Parent, 2011), TORC2 plays a key role in regulating the actin cytoskeleton in yeasts, *Dictyostelium discoideum* and mammalian cells (Jacinto *et al*, 2004; Baker *et al*, 2016; Lee *et al*, 2005). While the basic principle of control of each regulatory signal (e.g. phosphorylation and calcium signalling) are understood, the interplay between parallel modes of regulation is relatively unknown. *S. pombe*, contains both TORC1 and TORC2 complexes (Petersen, 2009).

In the current study, we have used molecular cell biological, biochemical and single molecule techniques to help identify and characterise a novel TORC2 phosphorylation-dependent system for regulating calcium-dependent switching of different calmodulin light chain(s) binding to the neck region of Myo1. We have established the contribution that each calmodulin plays in regulating this conserved motor protein and how they affect the conformation of the myosin lever arm. We propose a concerted mechanism of regulation by both calcium and phosphorylation that controls motility and function of Myo1 in response to signals controlling cell cycle progression.

## Results

### Fission yeast myosin-1 is phosphorylated within the IQ neck domain

Analysis of extracts from exponentially growing fission yeast cells indicates its sole class I myosin, Myo1, is subject to multiple phosphorylation events (Figure 1A). Phosphoproteomics studies (Carpy *et al*, 2014; Wilson-Grady *et al*, 2008) revealed a conserved phosphoserine residue located within the IQ motif containing neck region of class I & V myosins (Figure 1B). The location of this AGC family kinase consensus phosphoserine site (Pearce *et al*, 2010) has the potential to impact myosin activity and function by affecting conformation of the lever arm as well as light chain binding. A phosphospecific antibody was raised to confirm phosphorylation of the Myo1 serine 742 (Myo1^S742^), and established that it is phosphorylated in a TORC2 signalling and growth media dependent manner (Figure 1C-E). Consistent with the TORC2 dependent pathway modulating cell growth in response to media quality (Petersen & Nurse, 2007), replacing the serine with a non-phosphorylatable alanine residue within *myo1.S742A* cells resulted in an inability to inhibit growth when cultured in media containing minimal nitrogen (Figure 1F).

**Figure 1.**
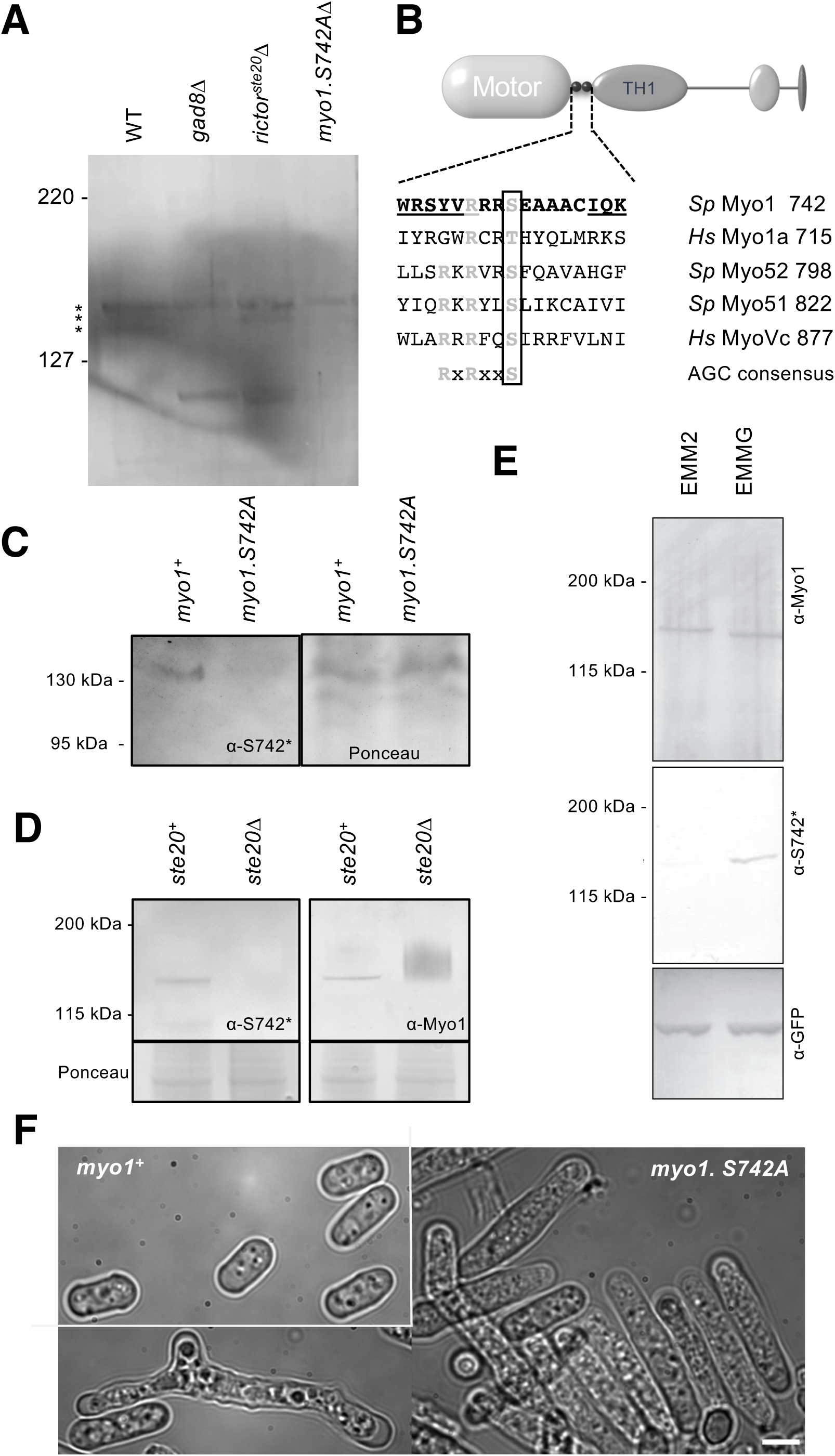
Myo1 serine 742 phosphorylation is TORC2 dependent. (A) Anti-Myo1 western blot of extracts from WT, *gad8*Δ*, ste20*Δ and *myo1.S742A* cells separated using Phos-Tag SDS-PAGE reveals Myo1 is subject to multiple phosphorylation events (*). (B) Sequence alignment of myosin IQ regions shows Myo1^S742^ lies within an AGC consensus sequence, conserved in class I and V myosins. (C) Western blots of extracts from *myo1*^+^ and *myo1-S742A* cells stained with Ponceau S and probed with phospho-specific anti-Myo1^S742^ antibodies demonstrate antigen specificity. (D) Myo1^S742^ is not phosphorylated in *ste20*Δ cells lacking the fission yeast TORC2 regulator Rictor^Ste20^. (E) Myo1^S742^ is phosphorylated in cells cultured in minimal media containing Glutamic acid (EMMG) but not in EMM2 with an ammonium chloride nitrogen source. (F) WT and *myo1.S742A* cells grown to starvation in EMMG for 72 hrs. In contrast to WT, *myo1.S742A* cells fail to stop growing upon media induced G1 arrest. Scale – 5 μm.

### Phosphorylation modulates Myo1 lever arm length

As serine 742 lies within the IQ motif containing neck region of myosin-1, we explored whether Myo1^S742^ phosphorylation affects calmodulin binding and conformation of the neck region. Isoforms of the Ca^2^+ sensitive fission yeast calmodulin (Cam1 and Cam1.T6C) were isolated in their native amino-terminally (Nt) acetylated forms using bacteria co-expressing the fission yeast NatA amino-α-acetyl-transferase complex (Eastwood *et al*, 2017). A FRET based fusion was generated with CyPet donor and YPet acceptor fluorophores (Nguyen & Daugherty, 2005) juxtaposed around the Cam1 protein to monitor Ca^2^+ dependent changes in Cam1 conformation (Figure 2A). This FRET-Cam1 fusion (Figure 2B), and Nt-acetylated IAANS labelled Cam1.T6C (Figure 2C) established Ca^2^+ binding brings about a change in the Cam1 conformation. Calculated pCa values for the Cam1-FRET (Figure 2B pCa_50_: 6.12), reflect global change in Cam1 conformation, while the IAANS dependent pCa (Figure 2C pCa_50_: 6.54) reflects Ca^2+^ dependent changes in the local environment at the amino lobe of Cam1. Quin-2, fluorescence of which increases upon Ca^2+^ binding (Tsien, 1980), was used to establish Ca^2+^ ions release from Cam1 with 3 distinct rate constants (137, 12.9 and 2.0 s^−1^) (Figure 2D).

**Figure 2.**
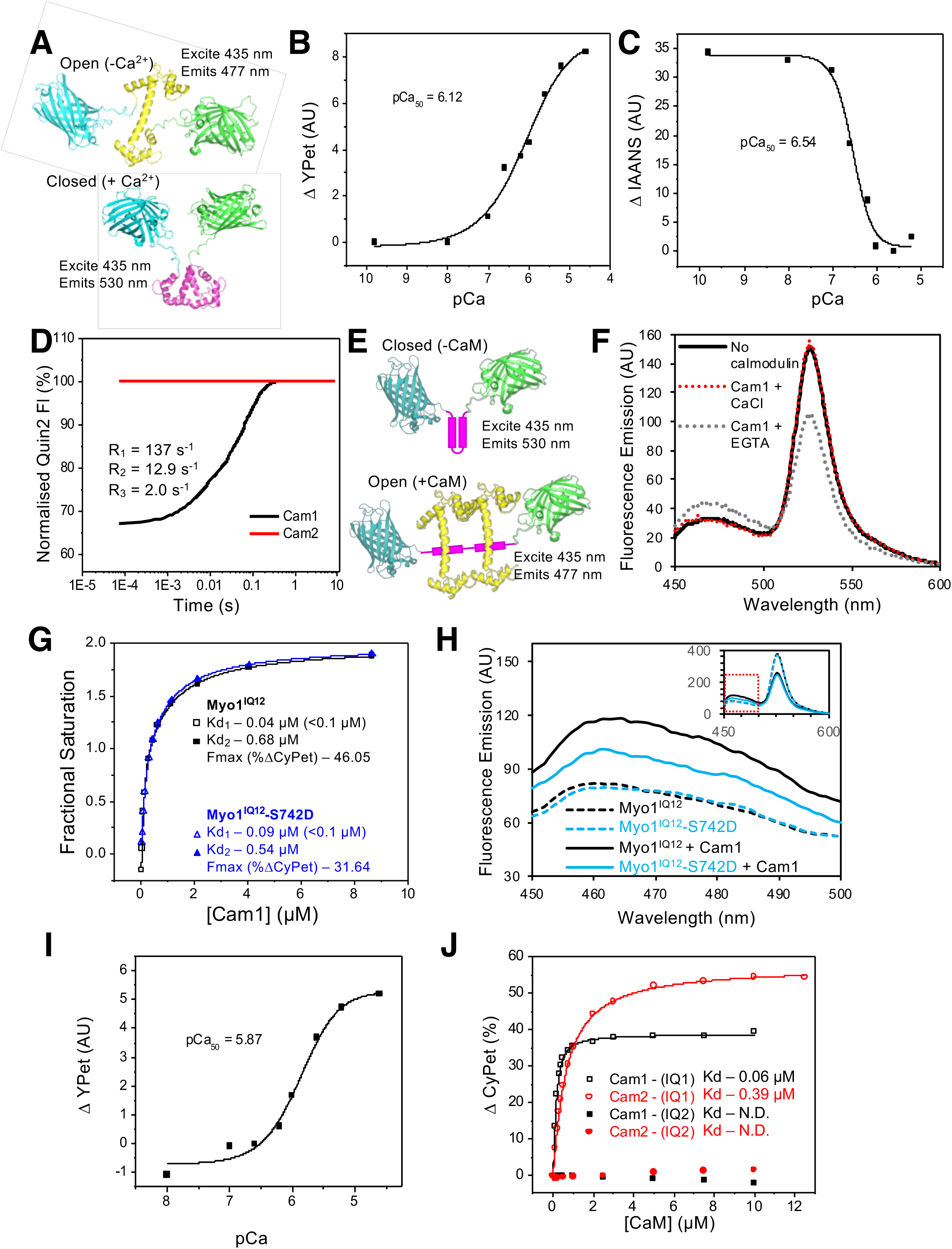
*In vitro* characterisation of interactions between Myo1 and Cam1. (A) Predicted models of the CyPet-Cam1-YPet FRET reporter protein (Cam1-FRET) in the absence (upper panel) and presence (lower panel) of Ca^2+^. (B) pCa curve plotting Ca^2+^ dependent changes of Cam1-FRET protein conformation (Δ in FRET signal). (C) pCa curve plotting Ca^2+^ dependent changes in IAANS fluorescence of IAANS labelled Cam1-T6C. (D) Transient curves of changes in Quin2 fluorescence brought by Ca^2+^ release from Cam1 (black) and Cam2 (red). (E) Predicted models of the CyPet-Myo1^IQ12^-YPet FRET reporter protein (Myo1IQ12-FRET) in the absence (upper panel) or presence (lower panel) of Calmodulin binding. (F) Spectra of Myo1^IQ12^-FRET reporter alone (black line) or with Cam1 in the presence Ca^2+^ (red dotted line) or absence (grey dotted line) of Ca^2+^. (G) Curves plotting Cam1 dependent changes of FRET donor signal of wild type (black) or S742D phosphomimetic (blue) Myo1^IQ12^-FRET proteins. (H) Spectra of Myo1^IQ12^-FRET (black traces) and Myo1^IQ12-S742D^-FRET (blue traces) in the absence (dashed lines) or presence (lines lines) of Cam1 illustrate differences in conformation of the Myo1 neck region. (I) pCa curve plotting Ca^2+^ dependent changes in acceptor fluorescence of Myo1^IQ12^-FRET. (J) Curves plotting Cam1 (black) and Cam2 (red) dependent changes of FRET donor signal of Myo1-FRET proteins containing single IQ domains (IQ1 – empty shapes; IQ2 – filled shapes).

To characterise Cam1 binding to the IQ neck region of the fission yeast myosin-1, recombinant FRET constructs were produced in which CyPet and YPet were separated by individual or both Myo1 IQ motifs (Myo1^IQ1^-FRET, Myo1^IQ2^-FRET, Myo1^IQ12^-FRET) (Figure 2E & S1). Cam1 binding to the IQ motif(s) stabilises the α-helix and results in a calcium regulated drop in FRET signal (Figure 2E-F). Analysis of interactions between Cam1 and Myo1^IQ12^-FRET revealed Cam1 molecules associated with the combined Myo1^IQ12^ motifs with 2 distinct phases, each contributing 50% of the overall change in signal (Figure 2G). The first Cam1-Myo1^IQ12^ binding event corresponds to an affinity of less than 0.1 μM (binding was too tight to calculate affinity with higher precision), while the second event correlates with an approximately 10-fold weaker binding affinity (0.68 μM). This association was seen to be sensitive to calcium (pCa of 5.87) (Figure 2I), illustrating Cam1 only associates with Myo1 in low cellular Ca^2+^ concentrations. Interestingly while Cam1 was seen to bind tightly to Myo1^IQ1^ alone (K_d_<0.1 μM), no detectable association was observed for the equivalent single Myo1^IQ2^ motif (Figure 2J). Together these data are consistent with a sequential cooperative binding mechanism by which the stable residency of Cam1 in the first IQ position is required before calmodulin can bind to Myo1^IQ2^.

Replacing serine 742 within the IQ neck region with a phosphomimetic aspartate residue had no significant impact upon the affinity, calcium sensitivity or cooperative nature of the interaction between Myo1 and Cam1 (Figure 2G). However, the phosphomimetic replacement resulted in a change in maximum FRET signal upon Cam1 binding (F_max_ 46.05 vs 31.64) (Figure 2G & H) indicating Myo1^S742^ phosphorylation changes the conformation of the lever arm upon Cam1 binding, rather than modulating the affinity for Cam1.

### Phosphorylation regulates Myo1 dynamics and endocytosis

To explore *in vivo* Myo1 and calmodulin dynamics we generated prototroph *S. pombe* strains in which endogenous *myo1, cam1, or cam2* genes were fused to cDNA encoding for monomeric fluorescent proteins (Figure 3A). Using high-speed (20 Hz) single molecule TIRF analysis we explored how Myo1^S742^ phosphorylation impacts Myo1 and Cam1 dynamics and function within the cell. Myo1 and Cam1 associated with the cell membrane in two distinct ways: we observed rapid transient associations of single molecules at the cell membrane, characterised by low-intensity single stepwise changes in intensity as well longer endocytic events which were much brighter and had a very different time-course. Single molecules of Myo1 and Cam1 bound transiently at the cell membrane and moved with low mobility (0.03 μm^2^.s^−1^), ∼10-times slower than diffusion of integral membrane proteins (Mashanov *et al*, 2010). The individual, diffraction-limited fluorescent spots appeared and disappeared in a stepwise fashion (i.e. within a single video frame). Event durations were exponentially distributed with mean lifetime of 2.2 s^−1^ (n = 152) (Movie 1). In contrast, during endocytic events, the fluorescence signal increased gradually, rising to a peak amplitude consistent with ∼45 molecules of mNeongreen.Myo1 (rate ∼13 molecules.s^−1^), which dwelled for ∼6 s, before falling to baseline (rate ∼14 molecules.s^−1^) (Figure 3B, Movie 2). The estimated number of Myo1 molecules is lower than reported in an earlier study (Sirotkin *et al*, 2010) perhaps due to differences in imaging techniques, as TIRF imaging illuminates the specimen to a depth of ∼100nm whereas confocal imaging would extend to > 400nm). The duration (Tdur) of endocytic events (measured as described in the Methods) was 13.84 s +/− 0.39 (mean +/− SEM, n=50) (Figure 3C) and while there was significant variation in the maximum mNeongreen.Myo1 intensity (2373 +/− 155), there was no correlation between maximum intensity and event duration (Figure 3D). Fluorescence intensity dynamics of Cam1.GFP during endocytic events were similar to mNeongreen.Myo1, but T_dur_ was significantly shorter (P <0.0001), 10.99 s +/− 0.21 (n=52) while the peak intensity was roughly double that measured for mNeongreen.Myo1 and equivalent to ∼ 90 GFP molecules (Figure 3E) consistent with Cam1 occupying both IQ sites within the Myo1 neck region. The briefer event duration observed for Cam1 might be explained by Cam1 dissociating from Myo1 before Myo1 leaves the endocytic patch. This idea was confirmed using two-colour imaging of *mNeongreen.myo1 cam1.mCherry* cells which showed Myo1 and Cam1 arrived simultaneously at the endocytic patch, but Cam1.mCherry disassociated ∼3 s before mNeongreen.Myo1 (Figure 3F, S2).

**Figure 3.**
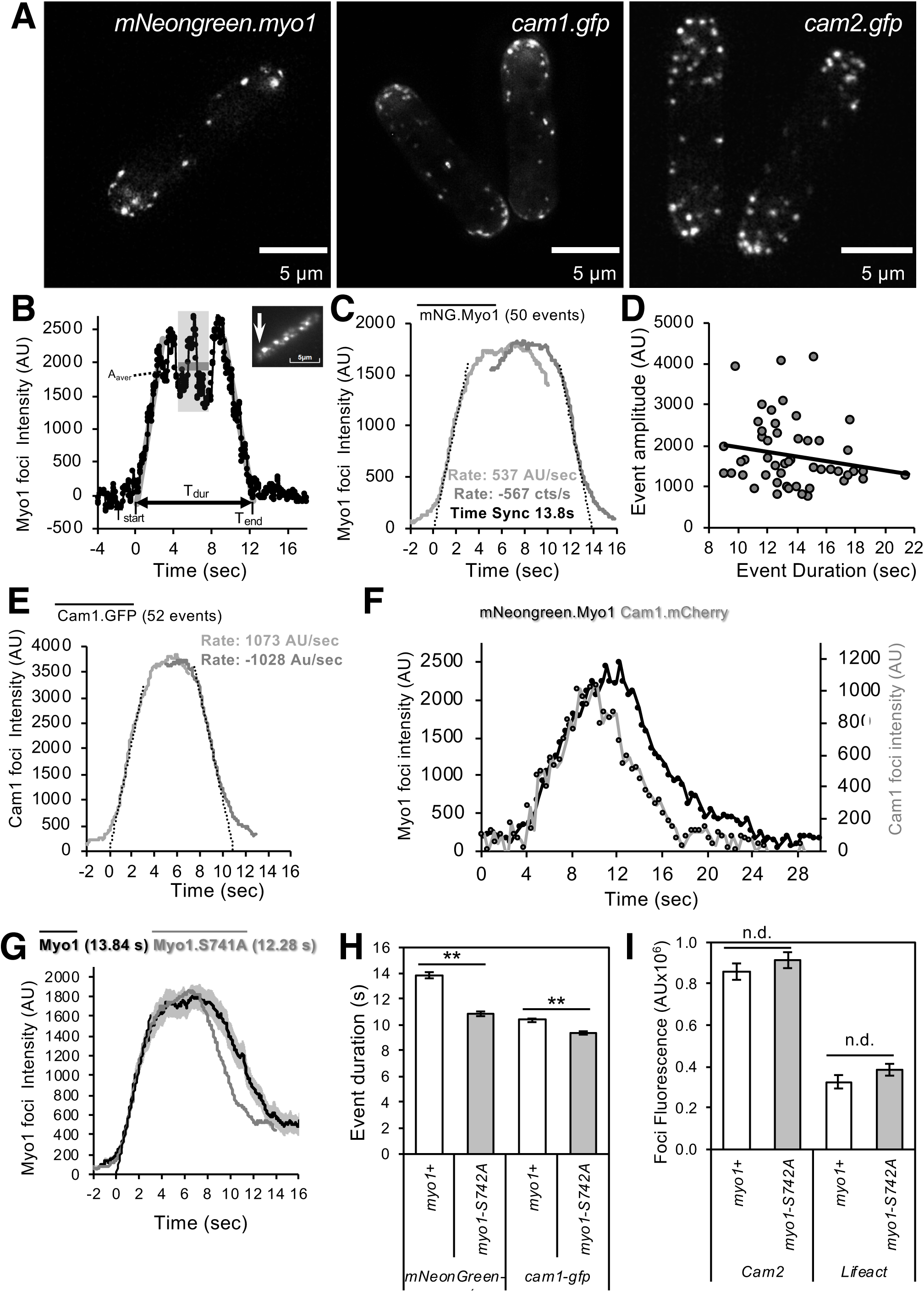
Myo1 and Cam1 dynamics in wild type and *myo1.S742A* cells. (A) Maximum projections of 31-z stack widefield images of *mNG.myo1, cam1.gfp* and *cam2.gfp* cells (Scales - 5 μm). (B) An example relative intensity trace of a mNeongreen.Myo1 endocytic event. Linear fitting (60 points) was used to find the highest slope for both rising and falling edges. The intercept with zero intensity level was used to calculate T_begin_, T_end_, and subsequently the duration of the event. Insert: An arrow shows the location of the endocytosis event (5X5 pixels area). (C) Averaged profile from 50 individual Myo1 membrane association events described in (B), synchronised relative to T_begin_ (grey line) and T_end_ (black line). (D) Plot of event duration (sec) against number of Myo1 molecules (fluorescence amplitude). (E) Averaged profile from 52 individual Cam1 membrane association events from TIRFM timelapse analysis of *cam1.gfp* cells. (F) Example of fluorescence trace from simultaneously tracking Myo1 and Cam1 membrane binding and disassociation events from TIRFM timelapse analysis of *mNeongreen.myo1 cam1.mCherry* cells. (G) Averaged profiles of combined averages of individual Myo1 (black line and grey s.d.) and Myo1.S742A (grey line) membrane association events from TIRFM timelapse analysis of *mNeongreen.myo1* and *mNeongreen.myo1.S742A* cells respectively. (H) Analysis of mean duration of Myo1 and Cam1 endocytic events in wt and *myo1.S742A* cells from widefield imaging (n > 30). Asterisks denote differences with >99% confidence. (I) Analysis of mean LifeACT and Cam2 signal at endocytic foci in WT and *myo1.S742A* cells (n > 30). No differences observed at 95% level of confidence. All error bars - s.d.

Analysis of Myo1 and Cam1 dynamics in *myo1.S742A* cells during endocytosis revealed Myo1^S742A^ had average assembly/disassembly rates and plateau intensity identical to wild type Myo1, but T_dur_ was 1.5 sec shorter (12.3s +/− 0.31 n=67) (Figure 3G & S2). Consistent with the *in vitro* data, the *myo1.S742A* mutation did not impact on the ability of Cam1 molecules associating at both IQ motifs, as average assembly/disassembly rates, and plateau intensity for Cam1 were the same in both wild type and *myo1.S742A* cells. However, we found that Myo1^S742A^ and Cam1 proteins disassociated simultaneously and somewhat earlier during the endocytic event in this strain.

These TIRF imaging data were consistent with widefield 3D-timelapse imaging that showed lifetimes of Myo1 and Cam1 foci were shorter in *myo1.S742A* cells when compared to *myo1*^+^ (Figure 3H). In contrast, while the *myo1.S742A* allele did not affect accumulation of Cam2 or LifeACT to sites of endocytosis (Figure 3I), the rate of endocytosis differs between old end and new ends of *myo1-S742A* cells compared to wild type (Figure 4A). Therefore, while Myo1^S742^ phosphorylation does not impact assembly of Myo1-Cam1 endocytic foci, it regulates myosin activity to change the function of the ensemble of endocytic proteins during bipolar growth.

**Figure 4.**
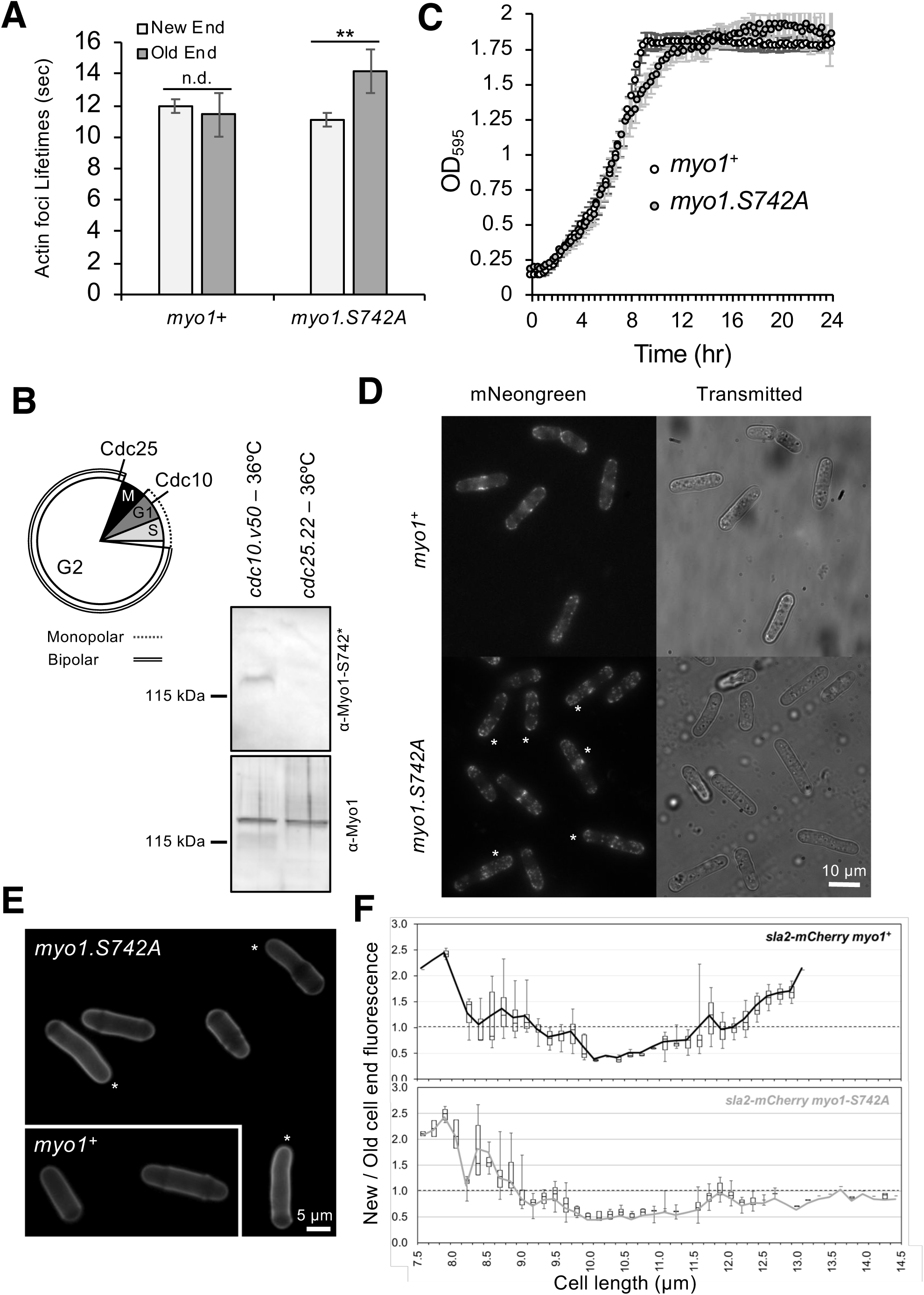
Myo1 S742 is phosphorylated in a cell cycle dependent manner to regulate polarised cell growth. (A) Actin foci periodicity at ends of WT and *myo1.S742A* cells in G2 phase (n>30). Asterisks denote difference with >99% confidence. (B) Graphic highlighting Cdc10 and Cdc25 execution points in relation to cell cycle phases and periods of monopolar / bipolar growth (left). Myo1^S742^ is phosphorylated in *cdc10.v50* arrested G_1_ cells, but not in pre-mitotic G_2_ *cdc25.22* arrested cells (right panels). (C) Averaged growth curves from 3 independent experiments of prototroph WT (empty circles) and *myo1.S742A* (grey filled circles) cells cultured in EMMG at 34 °C. Error bars - s.d. (D) Myosin-1 distribution and cell morphology of prototroph *mNeongreen.myo1*^+^ and *mNeongreen.myo1.S742A* cells cultured in EMMG at 34 °C. Asterisks highlight long bent cells. Scale - 10 μm. (E) Calcofluor stained WT and *myo1.S742A* cells. Asterisks highlight long bent cells displaying monopolar growth. Scale - 5 μm. (F) Ratio Sla2-mCherry fluorescence at “new”: “old” cell end, averaged from >30 growing mid-log *sla2-mCherry myo1*^+^ (upper panel) and *sla2-mCherry myo1.S742A* (lower panel) cells. Boxes plot median and quartile for each length measured, lines are plotted from the mean average value at each length measured.

### Myo1 S742 is phosphorylated in a cell cycle dependent manner to regulate polarised cell growth

Upon cell division fission yeast cells grow exclusively from the old cell end that existed in the parental cell. At a point during interphase (called New End Take Off-NETO) there is a transition to a bipolar growth (Mitchison & Nurse, 1985). This cell cycle switch in growth pattern correlates precisely with a parallel redistribution of endocytic actin patches (Marks & Hyams, 1985). As the *myo1.S742A* allele only affected actin dynamics at the old cell end during bipolar growth we examined whether this post-translational modification was subject to cell cycle dependent variance. Analysis of extracts from cell division cycle mutants arrested in G1 (*cdc10.v50* cells) or late G2 (*cdc25.22* cells) revealed Myo1^S742^ is phosphorylated in a cell cycle dependent manner (Figure 4B). This was confirmed by monitoring Myo1^S742^ phosphorylation in cells synchronised with respect to cell cycle progression (Figure S3). These data established that Myo1^S742^ phosphorylation peaks in early interphase (G1 cells), prior to the transition to a bipolar growth pattern, and steadily decreases until becoming undetectable towards the end of G2. Analysis of growth kinetics revealed *myo1.S742A* cells grow slower than wild type (Figure 4C), and have a longer average length (*myo1*^+^: 9.77 ± 1.77 μm; *myo1.S742A*: 13.2 ± 2.47 μm. t-test >99% significance n>500). In addition, a significant proportion of *myo1.S742A* cells demonstrate polarity defects, with 24.7% of cells having a bent morphology (i.e. growth deviates by >5° from longitudinal axis), compared to 1% seen in wild type (Figure 4D-E). Consistent with these observations, *myo1.S742* mutants exhibit defects in the transition from monopolar to polar growth. Cell wall staining revealed a significantly higher proportion of *myo1.S742A* cells exhibit monopolar growth compared to equivalent wild type, indicating disruption in the switch from monopolar to bipolar growth (Figure 4E). This was confirmed by tracking the cellular distribution of the actin patch marker, Sla2/End4, following cell division. Sla2 failed to redistribute to the newly divided end of *myo1.S742A* cells during interphase (Figure 4F). Together these data show that cell cycle variation in Myo1^S742^ phosphorylation modulates the myosin lever arm to regulate endocytosis and polarised growth.

### Cam2 associates with internalised endosomes and not Myo1 during vegetative growth

Myo1 has been reported to associate with a second calmodulin like protein, Cam2, via its second IQ motif (Sammons *et al*, 2011). However, our data indicate Cam1 occupies both Myo1 IQ motifs during endocytosis. Widefield microscopy revealed Myo1 and Cam1 dynamics (Figure 5A) at endocytic foci differ significantly from Cam2 which is recruited to sites of endocytosis later than Myo1 and Cam1, but prior to budding off, where, like CAPZA^Acp1^, Sla2 and actin, it remains associated with laterally oscillating internalised endosomes (Figure 5B-C). Similarly, simultaneous imaging of Cam1 and Cam2 in *cam1.mCherry cam2.gfp* cells revealed each protein localises to many foci lacking the other calmodulin, indicating differences in the timing of endocytic recruitment (Figure 5D). While Cam1 recruitment to endocytic foci is abolished in the absence of Myo1 (Figure 5E), the intensity, volume and number of Cam2 foci increases in the absence of Myo1 (Figure 5F Table 1). However, internalisation and lateral “oscillating” dynamics of Cam2, and actin were dependent on Myo1 (Figure 5F & G). Therefore, while Cam1 and Cam2 both localise to sites of endocytosis, they appear to do so at different times, and each have differing Myo1 dependencies.

**Figure 5.**
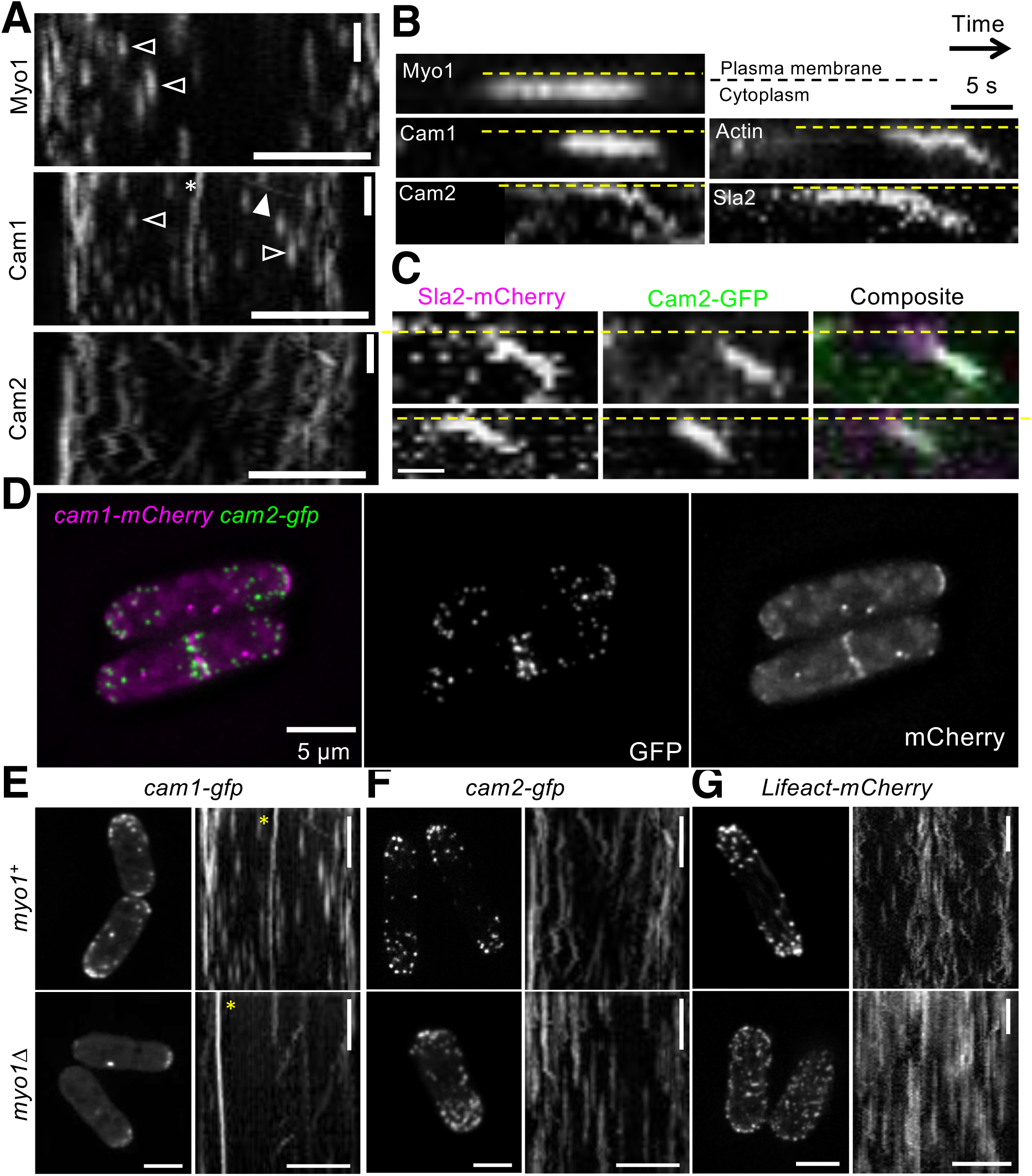
Cam2 associates with internalised endocytic vesicles. (A) Kymographs of GFP labelled foci from maximum projections of 13-z plane timelapse images of *mNeongreen.myo1* (upper panel), *cam1.gfp* (middle panel) and *cam2.gfp* (bottom panel) cells. Myo1 and Cam1 endocytic foci did not move on the membrane (black arrows). Spindle Pole Body (asterisk) and myosin V (white arrow) associated Cam1 are highlighted. In contrast Cam2 foci displayed extensive lateral movements. (B) Kymographs generated from single z-plane timelapse images of single endocytic foci surface during vesicle formation and subsequent internalisation. While Myo1 and Cam1 only associate with the plasma membrane, Cam2, Sla2 and actin are internalised on the vesicle after scission. (C) Kymographs of Cam2 and Sla2 co-internalisation in *sla2.mCherry cam2.gfp* cells. (D) Maximum projection of 31-z slice image of *cam1.mCherry cam2.gfp* cells reveals Cam1 (magenta) and Cam2 (green) colocalise to a subset of endocytic foci. (E-G) Single frames (left panels) and time kymographs (right panels) from maximum projections of 13-z plane timelapse images of *cam1.gfp* (E), *cam2.gfp* (F) and *LifeACT.mCherry* (G) in either *myo1*^+^ (upper panels) or *myo1*Δ (lower panels) cells show only Cam1 endocytic foci recruitment is dependent upon Myo1. Myo1 is required for internalisation of Cam2-GFP and LifeACT.mCherry labelled foci. Scales - 5 μm.

**Table 1:** AutoQuantX3 Image analysis data of wide-field fluorescence data of cells. Mutant strains were imaged in mix experiments with wild type cells, analysis for these cells is shown relative to the wild type control cells for each experiment. Statistical significance determined by an unpaired *t-test* is shown in brackets, a statistical significance of *p* < 0.05 is indicated in red.

| **Myo1** | ***mNeonGreen-myo1*** | ***mNeonGreen-myo1-S742A*** | ***YFP-myo1 cam2Δ*** | - |
| --- | --- | --- | --- | --- |
| Whole cell fluorescence (AU) | 9,453,813 | **0.86** (0.8628) | **0.90** (0.0295) | - |
| Cell size (μm<sup>2</sup>) | 98.9 | **0.84** (0.0863) | **0.95** (0.4542) | - |
| Maximum intensity (AU) | 33,477 | **0.92** (0.1446) | **0.62** (0.0001) | - |
| Number of foci | 15.9 | **0.82** (0.0203) | **0.84** (0.0261) | - |
| Average foci volume (μm<sup>3</sup>) | 0.98 | **0.98** (0.8595) | **0.64** (0.0001) | - |
| Total foci volume (μm<sup>3</sup>) | 15.2 | **0.75** (0.0020) | **0.53** (0.0001) | - |
| Total foci fluorescence (AU) | 139,712 | **0.76** (0.0061) | **0.45** (0.0001) | - |
| Average foci lifetime (s) | 14.0 | **10.9** (0.0001) | ND | - |
| N = | 32 | 30 | 37 | - |

| **Cam1** | ***cam1-gfp (myo1<sup>+</sup>)*** | ***cam1-gfp myo1-S742A*** | ***cam1-gfp cam2Δ*** | ***cam1-gfp myo1Δ*** |
| --- | --- | --- | --- | --- |
| Whole cell fluorescence (AU) | 61,530,900 | **0.89** (0.0197) | **0.97** (0.7145) | **1.14** (0.0733) |
| Cell size (μm<sup>2</sup>) | 86.1 | **0.99** (0.9271) | **1.08** (0.2016) | **1.00** (0.9748) |
| Maximum intensity (AU) | 251,700 | **0.82** (0.0563) | **0.72** (0.0001) | **1.41** (0.0001) |
| Number of foci | 14.1 | **0.96** (0.6960) | **0.93** (0.3200) | **0.42** (0.0001) |
| Average foci volume (μm<sup>3</sup>) | 1.12 | **0.67** (0.0081) | **0.74** (0.0321) | **1.52** (0.0188) |
| Total foci volume (μm<sup>3</sup>) | 14.33 | **0.68** (0.0004) | **0.78** (0.0703) | **0.56** (0.0001) |
| Total foci fluorescence (AU) | 1,020,350 | **0.63** (0.0002) | **0.66** (0.0013) | **0.60** (0.0001) |
| Average foci lifetime (s) | 10.4 | **9.4** (0.0001) | **13.3** (0.0001) | - |
| N = | 25 | 15 | 56 | 27 |

| **Cam2** | ***cam2-gfp (myo1<sup>+</sup>)*** | ***cam2-gfp myo1-S742A*** | - | ***cam2-gfp myo1Δ*** |
| --- | --- | --- | --- | --- |
| Whole cell fluorescence (AU) | 39,259,937 | **1.01** (0.8063) | - | **1.48** (0.0001) |
| Cell size (μm<sup>2</sup>) | 79.3 | **0.89** (0.2385) | - | **1.15** (0.3114) |
| Maximum intensity (AU) | 267,547 | **0.98** (0.6339) | - | **0.78** (0.0001) |
| Number of foci | 20.8 | **0.94** (0.4155) | - | **1.26** (0.0048) |
| Average foci volume (μm<sup>3</sup>) | 0.82 | **1.11** (0.0737) | - | **1.63** (0.0001) |
| Total foci volume (μm<sup>3</sup>) | 16.53 | **1.05** (0.4287) | - | **2.10** (0.0001) |
| Total foci fluorescence (AU) | 859,161 | **1.06** (0.3374) | - | **1.82** (0.0001) |
| N = | 20 | 31 | - | 17 |

Table 1:
| **LifeAct** | ***LifeAct (myo1<sup>+</sup>)*** | ***LifeAct myo1-S742A*** | ***ND*** | ***LifeAct myo1Δ*** |
| --- | --- | --- | --- | --- |
| Whole cell fluorescence (AU) | 17,116,300 | **1.14** (0.0936) | - | **1.15** (0.2851) |
| Cell size (μm<sup>2</sup>) | 84.5 | **1.01** (0.8787) | - | **1.14** (0.1529) |
| Maximum intensity (AU) | 94,671 | **1.06** (0.4403) | - | **0.64** (0.016) |
| Number of foci | 19.8 | **0.94** (0.5502) | - | **1.22** (0.0147) |
| Average foci volume (μm<sup>3</sup>) | 0.73 | **1.19** (0.0346) | - | **0.95** (0.6822) |
| Total foci volume (μm<sup>3</sup>) | 13.96 | **1.15** (0.1771) | - | **1.14** (0.2759) |
| Total foci fluorescence (AU) | 327,017 | **1.18** (0.1832) | - | **1.03** (0.8674) |
| N = | 23 | 23 | - | 23 |

TIRF analysis revealed on average a total of ∼30 Cam2 molecules recruit to each endocytic foci, and the kinetics of its recruitment to foci differ significantly to that observed for both Myo1 and Cam1. Cam2 often had a linear binding relationship (Figure 6A), which contrasts to the sigmoidal profiles observed for Myo1 and Cam1 (Figure 3C & E). TIRFM confirmed Cam2 remained associated with endocytic vesicles after they were internalised and their connection with the cell membrane was broken (Movie 3). Background corrected intensity traces of Cam2 dynamics at the membrane patch before, during, and after the end of endocytosis showed the signal rapidly dropped to baseline (<1s) (Figure 6A), with the Cam2 labelled vesicles remaining visible close to the membrane at the limit of the evanescent field. A large number of these mobile internalised Cam2 labelled vesicles were seen moving within the cytoplasm with relatively low cytosolic background signal (Movie 3), indicating much Cam2 associates with endocytic vesicles and remains bound to mature endosomes. During the latter stages of endocytosis, Cam2 was internalised on the endosome while Myo1 remained at the plasma membrane during endosome abscission (Sirotkin *et al*, 2010; Berro & Pollard, 2014; Picco *et al*, 2015). Timing of the Myo1 and Cam2 fluorescence signals did not correlate; Cam2 was associated with the endocytic vesicle moving away from the cell surface during endocytosis and remaining associated with the early endosome at the time of scission. Whereas, Myo1 and Cam1 remained immobile and stayed close to the cell surface (plasma membrane) throughout the endocytic cycle.

**Figure 6.**
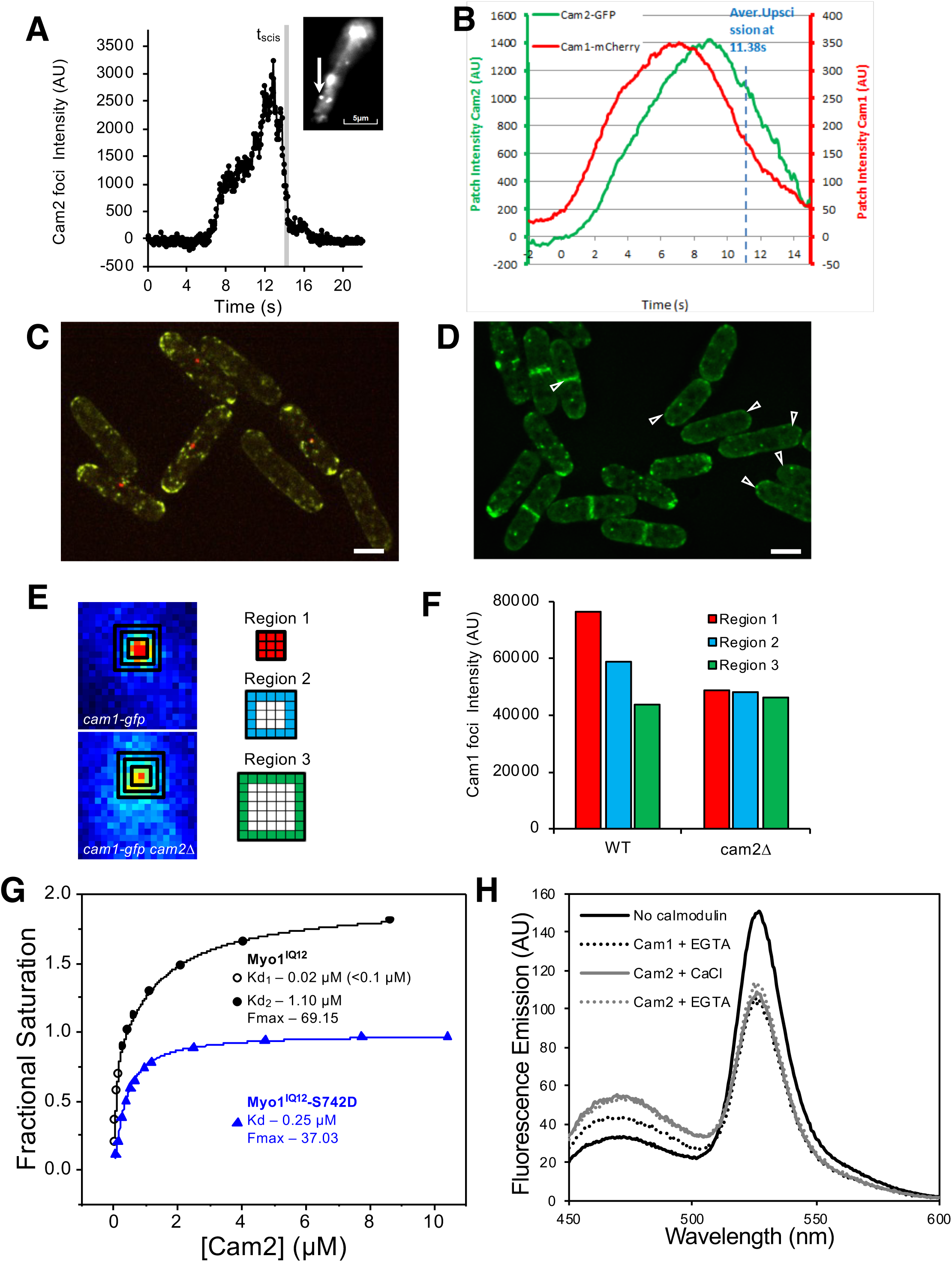
Cam2 impacts endosome organisation. (A) An example of the fluorescence trace of Cam2 membrane binding and vesicle internalisation event from TIRFM analysis of *cam2.gfp* cells. An abrupt drop in the fluorescence was marked as “scission time” (grey vertical line). An arrow shows the location of the monitored endocytic event (5X5 pixels area). (B) Averaged profile from 32 individual Cam2 membrane association events (green line) described in (A), together with Cam1-mCherry profile (red) from two-colour TIRFM imaging of *cam1.mCherry cam2.gfp* cells. Events were synchronized relative Cam1 T_begin_ Dashed line denotes mean timing of vesicle scission. (C) Maximum projection of 31-z slice widefield image of a mixture of prototroph *yfp.myo1 sid4.tdTomato* and *yfp.myo1 cam2*Δ cells. (D) Maximum projection of 31-z slice widefield image of a mixture of prototroph *cam1.gfp sid4.tdTomato* and *cam1.gfp cam2*Δ (arrows) cells. (E) Magnification of TIRF heat map of endocytic Cam1 in *cam2*^+^ (upper panel) and *cam2*Δ (lower panel) cells. Squares correspond to regions extending outward from centre of focus. (F) Plot of mean distribution of Cam1 across > 40 endocytic sites in WT and *cam2*Δ cells. (G) Curves plotting Cam2 dependent changes of FRET donor signal of wild type (black) or S742D phosphomimetic (blue) Myo1^IQ12^-FRET proteins. (H) Spectra of Myo1^IQ12^-FRET reporter alone (black line), with Cam2 in the presence Ca^2+^ (grey solid line) or absence (grey dotted line) of Ca^2+^, or with Cam1 in the absence of Ca^2^+ (black dotted line). Scales − 5 μm.

To correlate Myo1-Cam1 association at sites of endocytosis with scission of the endosome into the cytoplasm, we followed Cam1 and Cam2 dynamics simultaneously in *cam1.mCherry cam2.gfp* cells (Movie 4). An average curve generated from profiles of >30 complete individual endocytic events (Figure 6B) shows Cam2 moves away from the cell surface shortly after Cam1 leaves but before Myo1, with the time of abscission (Tscis) occurring on average 13.4 sec after the event starts (Tstart). Therefore endosome scission takes place during the Myo1 disassembly phase, and around the time Cam1 dissociates from Myo1.

Intriguingly, while the overall distribution of Myo1 and Cam1 appeared unaffected in *cam2*Δ cells, the number, volume and intensity of foci were significantly reduced (Figure 6C-D Table 1). TIRF-based analysis of the spatial distribution of Myo1 and Cam1 at endocytic foci revealed that Cam1 organised into more dispersed foci in the absence of Cam2 (Figure 6E-F), indicating Cam2 plays a role in organising the Myo1-Cam1 complex at the plasma membrane.

### Serine 742 phosphorylation increases the affinity of a single Cam2 for Myo1

*In vitro* analysis revealed two Cam2 molecules can associate with the unphosphorylated Myo1^IQ12^ region (Figure 6G) with 2 distinct phases. In contrast to Cam1 binding, 70% of the signal change is associated with an affinity of 1.10 μM. The smaller tighter signal change is not accurately measurable, but the combined change in signal is consistent with 2 binding events. As predicted from sequence analysis, Cam2 was not seen to associate with calcium (Figure 2D), and its conformation and interactions with Myo1 were insensitive to the divalent cation (Figure 6H). Like Cam1, Cam2 had a higher affinity for the first IQ motif (0.4 μM) than both IQ12 together, and did not bind to IQ2 alone (Figure 2J). Cam1 calcium binding, as measured by IAANS labelling or change in Quin-2 fluorescence were unaffected by Cam2, while gel filtration and fluorescence binding assays provided no evidence of a direct physical interaction between the two proteins (Figure S4). Interestingly a difference was observed in fluorescence amplitudes between Cam1 and Cam2 binding to the IQ12 motif, may indicate an impact upon lever arm length, (Figure 6H), potentially providing a mechanism to directly control Myo1 motor activity. Myo1^S742^ phosphorylation had no measurable impact upon the dynamics and distribution of Cam2 within fission yeast cells undergoing normal vegetative growth (Figure 7A Table 1). In contrast, *in vitro* analysis revealed Cam2 was only able to occupy one of the two IQ motifs in the Myo1^S742D-IQ12^ protein, be that with an increased affinity to the unphosphorylated protein (0.25 μM) (Figure 6G), indicating Cam2 impacts Myo1 function outside of the vegetative life cycle.

**Figure 7.**
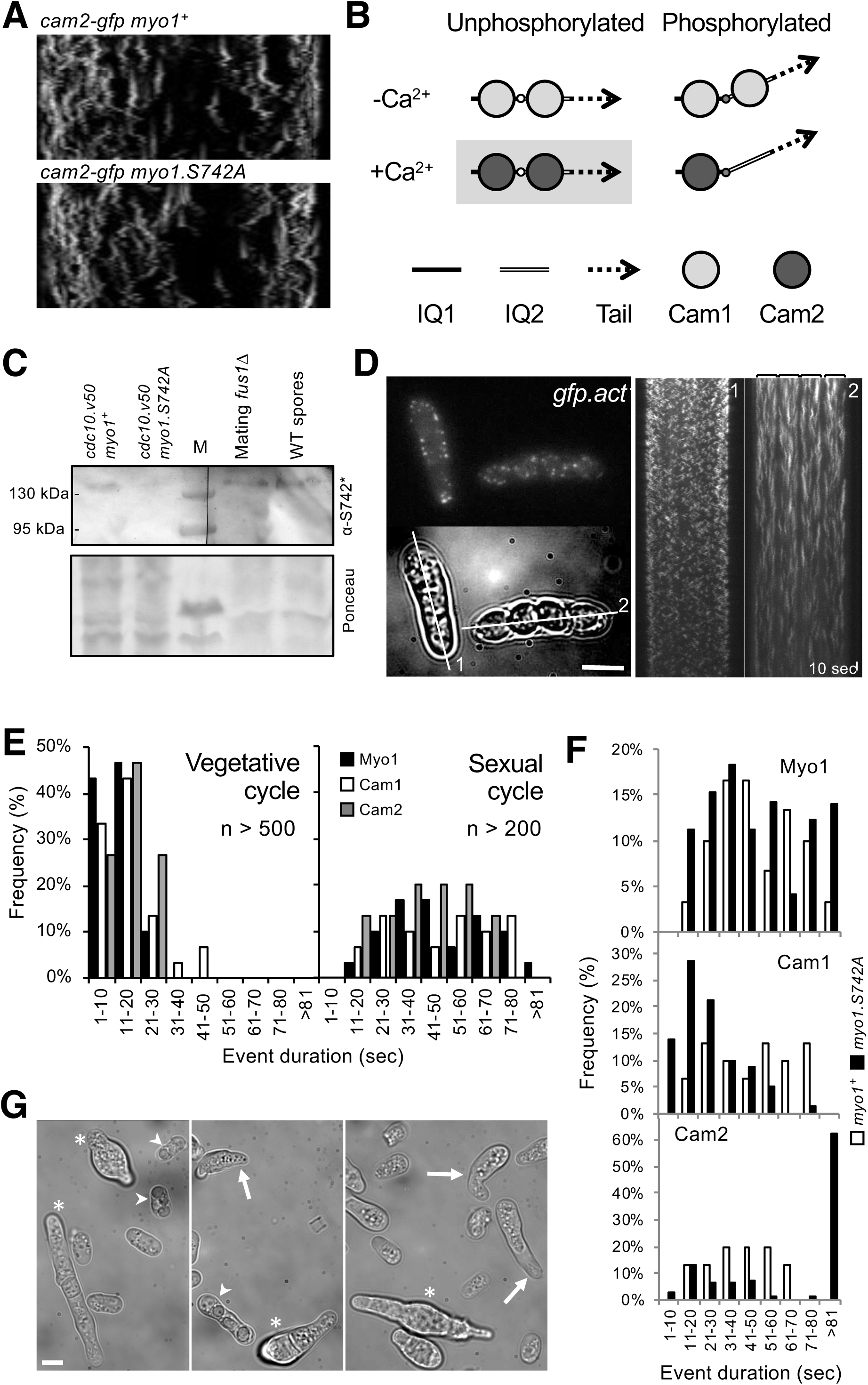
Myo1 S742 phosphorylation regulated Cam1 and Cam2 dynamics during meiosis. (A) Kymographs of Cam2.GFP foci dynamics in *myo1*^+^ (upper panel) and *myo1.S742A* (lower panel) cells. (B) Scheme of consequence of phosphorylation of Myo1 Ser742 (small empty circle) and Ca^2+^ levels upon Cam1 (light grey filled circle) and Cam2 (dark grey filled circle) binding to the IQ1 (solid thick black line) and IQ2 (compound line) motifs of Myo1, and impact on relative orientation of the myosin lever arm (dashed arrow). Highlighted combination of unphosphorylated Myo1^S742^ & Ca^2+^ does not normally occur in wild type cells. (C) Western blots of extracts from G1 arrested *cdc10.v50 myo1*^+^*, cdc10.v50 myo1-S742A* cells, conjugation arrested *fus1*Δ cells or spores, probed with phospho-specific anti-Myo1^S742^ antibodies confirm Myo1S742 remains phosphorylated throughout the sexual cycle. (D) Maximum projection of 13-z slice GFP fluorescence image and transmitted light image from a timelapse of vegetative (cell 1) and meiotic (cell 2) *gfp-act1* cells. Image from a GFP-act signal. Kymographs in the right panels were generated along the two dotted axes. (E) Histograms of lifetimes of Myo1 (black bars), Cam1 (white bars) and Cam2 (grey bars) foci in vegetative (left panel) and meiotic (right panel) cells. (F) Lifetimes of Myo1, Cam1 and Cam2 foci in WT (white bars) and *myo1.S742A* (black bars) meiotic cells. (G) Micrographs of *myo1.S742A* cell morphology on starvation media. * highlight cells with growth and polarity defects; arrows highlight cells with elongated or abnormally bent shmooing tips; and arrow heads highlight meioses resulting in defective spore formation. Scales − 5 μm.

### Cam1 and Cam2 associate with Myo1 during meiosis

Calcium levels within log phase yeast cells are relatively low (100–200 nM) (Ma *et al*, 2011; Miseta *et al*, 1999), and provides favourable conditions for Cam1 to associate with Myo1 (pCa - 5.87). Analysis of cell fluorescence indicated the relative abundance of Myo1 : Cam1 : Cam2 within the *S. pombe* cell to be 0.2 : 1.45 : 1 (Table 1), which is similar to the ratios defined by quantitative proteomic analysis of 0.45 : 1.56 : 1 (Marguerat *et al*, 2012). Similarly, image analysis of Cam1-GFP fluorescence revealed 1.7% of Cam1 to be associated with discrete foci within cells (Table 1), 40% of which is dependent upon Myo1, with the majority associating with the SPB (Figure 5D). This indicates ∼0.68% of cellular Cam1 associates with Myo1 at dynamic endocytic foci. These relative protein levels, binding affinities and low Ca^2+^ concentrations favour Cam1 binding to Myo1, over Cam2 at both IQ sites (Figure 7B), consistent with *in vivo* observations.

While Ca^2+^ levels are low during vegetative growth, sporadic prolonged calcium bursts occur upon pheromone release during mating (Carbó *et al*, 2016; Iida *et al*, 1990), and levels elevate significantly (∼10 fold) during the subsequent meiosis and sporulation (Suizu *et al*, 1995). Cam1 would be less likely to bind to Myo1 in these conditions (pCa 5.87). Myo1^S742^ is phosphorylated from G1, through cell fusion, persisting until completion of spore formation (Figure 7C). In addition Cam2 abundance increases significantly in relation to Cam1 during G1 upon mating and entry into meiosis (Mata & Bähler, 2006; Mata *et al*, 2002). These provide conditions that would favour Myo1-Cam2 interactions over Cam1 (Figure 7B), which is consistent with both Myo1 and Cam2 playing important role at the leading edge of forespore membrane formation during meiosis (Toya *et al*, 2001; Itadani *et al*, 2007). Consistent with this prediction, Myo1, Cam1, Cam2 foci lifetime and dynamics differ significantly to those observed in vegetative cells (P<0.0001), lasting significantly longer (>1 min) in meiotic and sporulating cells (Figure 7D & E). In contrast to vegetative cells, during meiosis and subsequent spore formation, like Myo1 and Cam1, Cam2 and actin foci were less dynamic, lacking any oscillation and remain in a fixed position with significantly longer lifetime than within actively growing cells (Figure 7D, Movie 5–8).

Finally, we used the *myo1.S742A* allele to explore the impact of Myo1^S742^ phosphorylation on Myo1, Cam1 and Cam2 dynamics and function during meiosis. In contrast to wild type, the lifetime of Cam1 foci were significantly shorter in *myo1.S742A* cells, and did not correlate with Myo1 and Cam2 dynamics, both of which differed significantly from *myo1*^+^ cells (Figure 7F). The majority of Cam2 foci remained present in the cell for greater than 2 mins in meiotic cells lacking Myo1^S742^ phosphorylation, which also differed significantly from Myo1^S742A^ dynamics, indicating normal Cam1 and Cam2 interactions with Myo1 were abolished. Consistent with *myo1.S742A* cells grown to stationary phase in minimal media (Figure 1F), heterothallic (h^90^) G1 arrested nitrogen starved *myo1.S742A* cells failed to inhibit polar growth (Figure 7G), mating cells accumulated with abnormal shmoo tips, and meiosis often resulted in cells with too few unequally sized spores (Figure 7G arrowheads). This spore defect phenotype is similar to that observed in *cam2*Δ cells (Itadani *et al*, 2007), which is consistent with a model whereby increase in cellular Ca^2+^ and Myo1^S742^ phosphorylation are both key for Cam2 association with and regulation of Myo1.

These data support a model by which changes in calcium levels and TORC2 dependent phosphorylation status provide a simple two stage mechanism for modulating motor activity by modifying lever arm length as well as switching calmodulin light chain preference to regulate myosin function in response to changing needs of the cell (Figure 7B).

## Discussion

Myosins are subject to diverse systems of regulation, which include composition of the actin track, cargo and light chain interactions, as well as phosphorylation. Here we describe a newly discovered mechanism by which phosphorylation of the myosin heavy chain (Figure 1) regulates light chain specificity, lever arm conformation and flexibility, to modulate and control cellular function. During the vegetative life cycle, within basal levels of cellular calcium, the fission yeast myosin-1 preferentially associates with two molecules of the calcium regulated calmodulin light chain Cam1 (Figures 2 & 3). During early stages of the cell cycle TORC2 dependent phosphorylation of the Myo1 neck region, to which the light chain(s) bind, changes the length of the Cam1 associated lever arm to moderate its activity to regulate the rate of endocytosis (Figure 4).

During the sexual cycle, Myo1^S742^ remains phosphorylated (Figure 7). This combined with the increase in cytosolic Ca^2+^ levels leads to a switch in light chain preference to a single molecule of the calcium insensitive calmodulin like, Cam2. The single Cam2 molecule is likely to bind IQ1 of S742 phosphorylated Myo1, as comparison with the structure of the IQ region of Myosin-1 and calmodulin (Lu *et al*, 2014), phosphorylation of S742 is likely to impact calmodulin interactions at the 1^st^ IQ position. Furthermore, our data reveals that Cam2 is unable to associate with IQ2 alone, as it is necessary for one calmodulin to occupy IQ1 in order for a second to bind to IQ2. This switch in light chain occupancy may provide a mechanism to change the stiffness of the Myo1 neck region (i.e. the “lever arm”) and thereby modulate the movement and force it produces during the acto-myosin ATPase cycle and/or the load-sensitivity of its actin-bound lifetime.

While Myo1 is capable of associating with phospholipid membranes via its Plekstrin Homology (PH) domain, *in vivo* data suggests that this alone is not sufficient to enable a stable interaction at the plasma membrane (Figure 8A). The build-up of the early endocytic markers, such as Pan1 or Sla1, are necessary to catalyse its nucleation to early endocytic patches allowing Myo1 foci to form at the site of membrane invagination. This is consistent with our observation that once initiated, Myo1-Cam1 foci do not collapse, but go on to complete the endocytic event (Figure 8B) (Sun *et al*, 2015; Barker *et al*, 2007). Similarly, the size of this early marker “patch” has a direct impact upon the number of Myo1 molecules recruited to the plasma membrane, which is consistent with the role of Pan1 in enhancing the Arp2/3 actin nucleating activity of myosin-1 foci in yeast (Barker *et al*, 2007).

**Figure 8.**
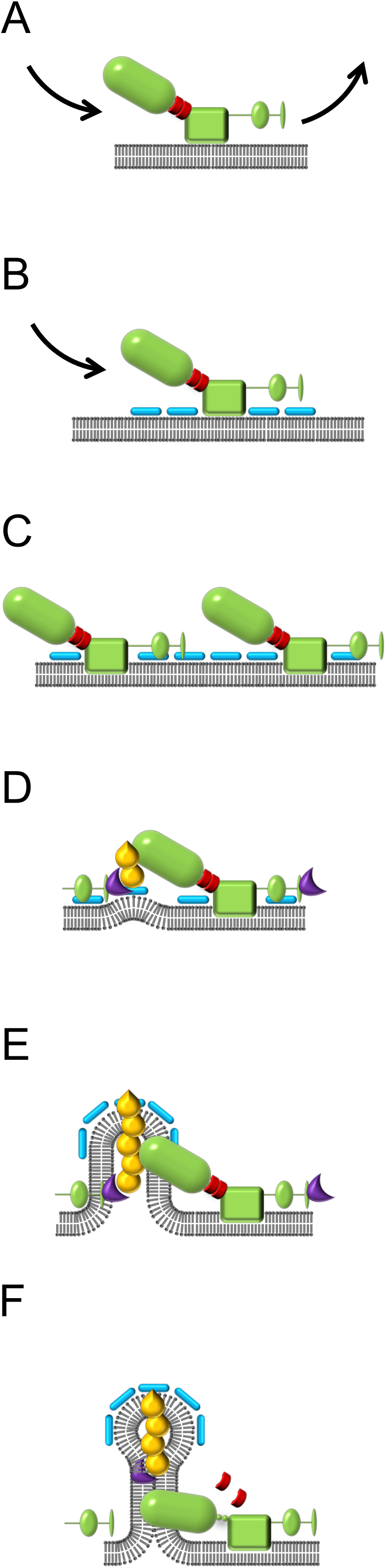
Model of Myo1 tension dependent interactions at the plasma membrane. (A) Myo1 (green) transiently associates with the plasma membrane. (B) In the presence of early markers of endocytosis (blue) this interaction is stabilised, and Myo1 accumulates to a critical concentration at the endocytic foci (C), whereupon myosin heads associate with growing Arp2/3 (purple) nucleated actin polymers (yellow) attached to the membrane (D), and monitor tension between the actin filament and internalised plasma membrane (E). Upon release of the calmodulin light chain (red), the myosin-1 would its ability to monitor tension and subsequently disengage from the actin polymer and membrane (F).

The local concentration of Myo1 at the endocytic patch appears critical, rather than the absolute number of Myo1 molecules, as the latter does not affect the duration of the Myo1 driven event. Indeed the duration of Myo1’s residency at the plasma membrane is driven by Cam1 and phosphorylation regulated neck length. Interestingly neither of these factors affect the rate of Myo1 or Cam1 recruitment or disassociation from the membrane.

Therefore the size of the Pan1 patch determines the number of Myo1 molecules necessary to generate a critical local concentration of Arp2/3 nucleated actin filaments (Figure 8C) (Barker *et al*, 2007). At the critical concentration myosin heads are able to interact with actin filaments nucleated from either adjacent Myo1 tails or WASP activated Arp2/3 complexes, tethered to the membrane via molecules such as the Talin like Sla2 (Figure 8D) (Sirotkin *et al*, 2005; 2010). The Myo1 is then primed to act as a tension sensor against the actin filament, as it pushes against the membrane of the internalised endosome, which grows against the significant 0.85 MPa (8.3 atm) turgor pressure within the cell (Minc *et al*, 2009) (Figure 8E). While observations within budding yeast indicate motor activity from a ring of myosins at the lip of the endosome (Mund *et al*, 2018) is necessary for endocytic internalisation the mechanism by which the myosin interacts with actin to facilitate this is unknown (Sun *et al*, 2006).

The number of Myo1 molecules at the plasma membrane foci remains constant, as the membrane is internalised, until 2 seconds after Cam1 disassociates from Myo1 (Figure 8F). While the trigger for Cam1 release is unknown, the rapid ensemble nature of the event indicates it is likely to be initiated by a rapid localised spike in calcium. This could perhaps be driven by a critical level of membrane deformation coupled to calcium influx - similar to processes proposed for mechano-transduction and the role of mammalian myosin-1 within the stereocilia of the inner ear (Adamek *et al*, 2008; Batters *et al*, 2004). Alternatively, mechanical forces acting on Myo1 may drive Cam1 dissociation. Genetics studies from budding yeast indicate that calmodulin mutants, unable to bind Ca^2+^, release normally from myosin-1 (Geiser *et al*, 1991). Research using mammalian brush border myosin-1, indicates that changes in lipid composition of membranes to which the motor is associated are sufficient to displace calmodulin from the IQ region (Hayden *et al*, 1990). In fission yeast this change in lipid could be rapidly triggered by PI4-kinase phosphorylation (Cam2 is the light chain for PI4 kinase (Sammons *et al*, 2011)). This is consistent with timing of Cam2 membrane recruitment and could go some way to explain why Myo1 foci are more dispersed in absence of Cam2.

Once Cam1 detaches from the Myo1 molecule, the neck loses rigidity (Figure 8F), reducing tension between the myosin motor and actin filament, causing it to detach rapidly from F-actin (Lewis *et al*, 2012; Mentes *et al*, 2018). Given the off-rate of single Myo1 molecules from the plasma membrane is ∼2 sec^−1^ (Figure 3B), lack of association with actin would mean that Myo1 would leave the endocytic patch a second or so after losing its Cam1 light chain. Together these events account for the 2 sec delay between disappearance of Cam1 and Myo1 from the membrane. The same drop in tension at the plasma membrane could provide the signal for scission of the endosome (Palmer *et al*, 2015).

The conformation and rigidity of the Myo1 lever arm would therefore play a key role in modulating the tension sensing properties of the motor domain. This is consistent with our data, where wild type phosphorylatable Myo1 resides at the membrane ∼1.8 sec longer than unphosphorylated Myo1^S742A^ (Figure S2). Phosphorylation-dependent changes in the conformation of the myosin neck provide a simple mechanism to modulate the rate of endocytosis according to the size and needs of the cell. Similarly, in the presence of Ca^2+^ and Myo1^S742^ phosphorylation, a single Cam2 resides at IQ1 motif of the neck (Figure 7B), again modulating neck conformation adjacent the motor domain as well as allowing flexibility within the carboxyl half of the neck region. This would provide a relatively tension insensitive motor, that stalls against the actin polymer, and would therefore persist significantly longer at the endocytic foci, as observed here (Figure 7E). These changes in lever arm properties change the overall rate of endocytosis, as observed in differences for actin labelled endosomes to internalise (Figure 4A).

Thus phosphorylation-dependent changes in the calcium regulated conformation and rigidity of the myosin lever arm could provide a universal mechanism for regulating the diverse cytoplasmic activities and functions of myosin motors within all cells.

## Materials and Methods

### Yeast cell culture

Cell culture and maintenance were carried out according to (Moreno *et al*, 1991) using Edinburgh minimal medium with Glutamic acid nitrogen source (EMMG) unless specified otherwise. Cells were cultured at 25 °C unless stated otherwise and cells were maintained as early to mid-log phase cultures for 48 hours before being used for analyses. Genetic crosses were undertaken on MSA plates (Egel *et al*, 1994). All strains used in this study were prototroph and listed in Supplementary Table 1.

### Molecular Biology

*cam1*^+^ (SPAC3A12.14), *cam1.T6C* and *cam2*^+^ (SPAC29A4.05) genes were amplified as *Nde1 - BamH1* fragments from genomic *S. pombe* DNA using o226/o227 and o393/o394 primers and cloned into pGEM-T-Easy (*Promega*, Madison, WI, USA). After sequencing the subsequent genes were cloned into pJC20 (Clos et al., 1990) to generate bacterial calmodulin expression constructs. DNA encoding for the FRET optimized fluorophores CyPet and YPet (Nguyen and Daugherty, 2005) were each amplified using primers o405 / o406 and o403 / o404 respectively. o406 also incorporated DNA at the 3’ end of the CyPet ORF encoding for the first IQ motif of the Myo1 neck region, while o404 included DNA encoding a Gly3His6 tag at the 3’ of the YPet ORF. The two DNA fragments were cloned into pGEM-T-Easy in a three-way ligation reaction to generate pGEM-CyPet-Myo1IQ1-YPet. The CyPet-Myo1^IQ1^-YPet DNA was subsequently sequencing and cloned as a Nde1 - BamH1 fragment into pJC20 (Clos & Brandau, 1994) to generate pJC20CyPet-Myo1^IQ1^-YPet. Complementary oligonucleotides o425 & o426 were annealed together and ligated into BglII – Xho1 cut pJC20CyPet-Myo1^IQ1^-YPet to generate pJC20CyPet-Myo1^IQ12^-YPet. Similarly, complementary oligonucleotides o429 & o430 were annealed together and subsequently ligated into Sal1-BglII cut pJC20CyPet-Myo1^IQ1^-YPet and the subsequent Xho1 fragment was excised to generate pJC20CyPet-Myo1^IQ2^-YPet. Site directed mutagenesis was carried out using pJC20CyPet-Myo1^IQ12^-YPet template and o427 & o428 primers to generate pJC20CyPet-Myo1^IQ12^S742D-YPet. Complementary oligonucleotides o449 & o450 were annealed together and ligated into Nru1 – Xho1 digested pJC20CyPet-Myo1^IQ12^S742D-YPet to generate pJC20CyPet-Myo1^IQ12^S742A-YPet. All plasmids were sequenced upon construction. Strains with fluorophore tagged alleles of *cam1*^+^ and *cam2*^+^ were generated as described previously using appropriate template and primers (Bähler *et al*, 1998). Strains in which the *myo1.S742A, myo1.S742D, mNeongreen-myo1*, *mNeongreen-myo1.S742A*, or *mNeongreen-myo1.S742D* alleles replaced the endogenous *myo1*^+^ gene (SPBC146.13c) were generated using a marker switching method (MacIver *et al*, 2003). Oligonucleotides are described in Supplementary Table 2.

### Protein expression & purification

All recombinant proteins were expressed and purified from BL21 DE3 *E. coli* cells, except Cam1 proteins where BL21 DE3 pNatA cells (Eastwood *et al*, 2017) were used to allow amino-terminal acetylation (Figure S1). *Calmodulin purification*: Cell lysates were resuspended in Buffer A (50 mM Tris, 2 mM EDTA, 1 mM DTT, 0.1 mM PMSF, pH 7.5) and precleared by high speed centrifugation (48,500 RCF; 30 min; 4 °C), before ammonium sulphate was added to the supernatant at 35 % saturation, incubated for 30 minutes at 4 °C. Precipitated proteins were removed by centrifugation (48,500 RCF; 30 min; 4 °C). For Cam1 purifications the precipitation cleared supernatant was added to a pre-equilibrated 10 ml phenyl sepharose (CL-4B) column (Buffer B: 50 mM Tris, 1 mM DTT, 1 mM NaN_3_, 5 mM CaCl_2_, pH 8.0), washed in 4 volumes of Buffer B before eluted as fractions in Buffer C (50 mM Tris, 1 mM DTT, 1 mM NaN_3_, 5 mM EGTA, pH 8.0). For Cam2 purification the precipitation cleared supernatant underwent a second round of ammonium sulphate precipitation and clearing, and the subsequent supernatant subjected to isoelectric precipitation (pH 4.3) and centrifugation (48,500 RCF: 30 minutes; 4 °C). The resultant pellet was resuspended in Buffer A, heated to 80 °C for 5 minutes and denatured proteins removed by centrifugation (16,000 RCF; 5 min). *His-tagged* proteins were purified in native conditions using prepacked, pre-equilibrated 5ml Ni^2+^ columns.

### Fast reaction kinetics

All transient kinetics were carried out using a HiTech Scientific DF-61 DX2 Stopped Flow apparatus (TgK Scientific, Bradford-upon-Avon, UK) at 20°C. All data was acquired as the average of 3–5 consecutive shots and analysed using the KineticStudio software supplied with the equipment. Quin-2 fluorescence was excited at 333 nm and used a Schott GG445 cut off filter to monitor fluorescence above 445 nm. IAANS (2-(4’-(iodoacetamido)anilino)-naphthalene-6-sulfonic acid) was excited at 335 nm and fluorescence was monitored through a GG455 filter. For the FRET measurements, CyPet was excited at 435 nm and YPet emission was monitored through a combination of a Wrattan Gelatin No12 (Kodak) with a Schott GG495 nm filter to monitor fluorescence at 525–530 nm.

### Fluorescence spectra

Emission spectra were obtained using a Varian Cary Eclipse Fluorescence Spectrophotometer (Agilent Technologies, Santa Clara, CA) using a 100 μl Quartz cuvette. For FRET measurements samples were excited at 435 nm (CyPet excitation) and emission was monitored from 450 – 600 nm with both slits set to 1 nm. Affinity experiments were carried out using 1 μM IQ-FRET protein with varying concentrations of Cam1 or Cam2 in a final volume of 100 μl in analysis buffer of 140 mM KCl, 20 mM MOPS, pH 7.0 with or without 2 mM MgCl_2_ and with 2 mM of EGTA, CaCl_2_ or Ca^2+^-EGTA as required.

### Live cell imaging

Live cell widefield fluorescence imaging was undertaken as described previously (Baker *et al*, 2016). For Total Internal Reflection Fluorescence Microscopy (TIRFM) *S. pombe* cells were immobilized on N<u>o</u>1, Ø 25 mm lectin coated coverslips and placed into imaging chambers filled with EMMG medium. A previously described custom TIRF Microscope (Mashanov *et al*, 2003) was used to image individual cells at a rate of 20 fps in either single of dual colour mode. Lasers: 488 nm/100 mW and 561 nm/150 mW (*Omicron*, Germany); emission filters 525/50 nm and 585/29 nm, dichroic mirror 552 nm (*Semrock*, NY); all lenses and mirrors (*Thorlabs*, NJ), except two Ø 3 mm mirrors (*Comar Optics*, UK) which directed light in and out of the 100 × 1.45 NA objective lens (*Olympus*, Japan). Sequences of images were captured using one or two iXon897BV cameras (*Andor Technology*, UK) with custom made acquisition software. 100% laser power (488 nm) was used to image individual mNeongreen-Myo1 and Cam1-GFP molecules. The laser intensity was reduced to ≤ 20% during endocytosis imaging experiments to minimize photobleaching. All imaging was undertaken at 23 °C.

### Image analysis

*Widefield data* was analysed using Autoquant software (*MediaCybernetics*, Rockville, MD, USA). All 3d image stacks were subjected to blind 3d deconvolution before analysis. Average size and number and cellular distribution of foci were calculated from all foci present within ≥ 30 cells for each sample examined. Timing of foci events were calculated from kymographs generated in Metamorph software (*Molecular Devices*, Sunnyvale, CA, USA). The proportion of cells displaying a bent cell phenotype was determined from more than >350 calcofluor (1 mg.ml^−1^) stained cells for each strain. Bent cells were defined by a deviation in the direction of growth of > 5° from the longitudinal axis. *TIRF data* analyses, including single molecule detection and tracking, was undertaken using GMimPro software (Mashanov & Molloy, 2007). Endocytic events were identified by creating an image representing the standard deviation of each pixel over the whole video sequence (known as a “z-projection”). Bright spots in this image correspond to regions of the yeast cell that showed large intensity fluctuations. Regions of interest (ROIs) ∼ 0.5 μm diameter (5×5 pixels) were created to enclose the site of endocytosis and changes in the averaged ROI intensity over the entire video record were saved for future analysis. To correct for local variation in background signal, the average intensity in a region 1.5 μm diameter around the endocytosis site (but not including the central ROI) was subtracted. Data from ROIs that were contaminated by other endocytosis events, occurring in close proximity and close in time, were manually excluded from the analysis. It was critical to identify accurately the start and end of each endocytosis event so that individual traces could be averaged. To facilitate this, the rising and falling phases of the intensity trace were fitted with a straight line (60 data points, 3 sec duration), see Figure 3C for example. The intercept of this line with the baseline intensity gave the t_start_ and t_end_ values and event duration (T_dur_ = t_end_ - t_start_) (see Figure 6A). Intensity traces for each given condition were synchronised to the starting point (t_start_) and averaged (except Cam2-GFP traces which were synchronised using t_start_ measured from simultaneously acquired Cam1-mCherry signal). Similarly, traces were synchronised to their end point (t_end_) and averaged. The mean duration of the events (T_dur_) for each condition was then used to reconstruct the mean intensity changes with calculated errors for event amplitude and timing (Table 2). Since the falling and rising phases of most events fitted well to a simple linear equation, the *slope* of the fitted lines was used to estimate the rate of accumulation and dissociation of the fluorescent molecules. As Cam2-GFP remained bound to the endocytic vesicle, when vesicle scission occurred intensity fell rapidly to zero as the vesicle diffused from the TIRF evanescent field; the time of scission was defined as t_scis_ (Figure 6C). Single particle tracking was performed using, GMimPro (Mashanov & Molloy, 2007) (ASPT module) so that the paths (or trajectories) of individual Myo1 molecules bound to cell membrane could be traced. Trajectories were analysed to yield mean intensities for individual NeonGreen and eGFP labelled proteins, which could be used to estimate the number of fluorescently-tagged molecules associated with each endocytotic event. Intensity-versus-time plots were generated from averages of >30 foci for each protein in each genetic background examined.

**Table 2:** Image analysis data of TIRF data of cells of the indicated genotype.

| Protein (*myo1 allele*) | Duration (SEM) | Amplitude (SEM) | Rise rate (SEM) | Drop rate (SEM) | N |
| --- | --- | --- | --- | --- | --- |
| Myo1 (*myo1*<sup>+</sup>) | 13.84(0.39)<sup>1,2,3</sup> | 2373(155) | 536(40.4) | 567(43) | 50 |
| Cam1 (*myo1*<sup>+</sup>) | 10.99(0.21)<sup>1</sup> | 4539(292) | 1074(83) | 1028(69) | 52 |
| Myo1 (*myo1.S742A*) | 12.28(0.31)<sup>2,4</sup> | 2274(128) | 536(34) | 570(41) | 67 |
| Cam1 (*myo1.S742A*) | 12.15(0.38)<sup>3,4</sup> | 4629(301) | 1153(98) | 1031(77) | 43 |

Significance differences observed in durations of: <sup>1</sup> Myo1 (wt cells) and Cam1 (wt cells) foci $p < 0.0001$ ; <sup>2</sup> Myo1 (wt cells) and Myo1 (*myo1.S742A* cells) foci $p < 0.002$ ; <sup>3</sup> Myo1 (wt cells) and Cam1 (*myo1.S742A* cells) foci $p < 0.0064$ .
<sup>4</sup> No significant difference observed between duration of Myo1 (*myo1.S742A* cells) and Cam1 (*myo1.S742A* cells) foci $p < 0.79$ .

## Supporting information

## Acknowledgements

We thank Professors M. Balasubramanian, I. Hagan, P. Nurse, C. Shimoda and T. Pollard for strains; and Dr Ben Goult for stimulating discussions and comments on the manuscript. This work was supported by the University of Kent and funding from the Biotechnology and Biological Sciences Research Council (BB/J012793/1 & BB/M015130/1), a Royal Society Industry Fellowship to DPM; a CASE industrial bursary from Cairn Research Ltd to KB and by the Francis Crick Institute which receives core funding from Cancer Research UK (FC001119), the UK Medical Research Council (FC001119) and the Wellcome Trust (FC001119) GIM and JEM.

## Supplementary Data Legends

**Supplementary Figure 1. Purified proteins used during *in vitro* studies.** Coomassie stained SDS-PAGE gel of recombinant proteins expressed and purified during this study. From left to right lanes contain (L) protein standard; (1) Nt-acetylated Cam1; (2) Nt-acetylated Cam1-T6C; (3) Cam1-FRET; (4) Cam2; (5) IQ12 peptide (not used during this study); (6) Myo1IQ12-FRET; and (7) Myo1IQ12S742D-FRET.

**Supplementary Figure 2. Relative TIRF profiles.** Combined profiles of averages from TIRFM timelapse analysis of Myo1 and Cam1 dynamics in wild type or *myo1.S742A* strains. (A) Myo1 (blue) and Cam1 (red) membrane association in wild type cells. (B) Myo1 membrane association in wild type (blue) and *myo1.S742A* (red) cells. (C) Myo1 (blue) and Cam1 (red) membrane association in *myo1.S742A* cells. (D) Cam1 membrane association in wild type (blue) and *myo1.S742A* (red) cells.

**Supplementary Figure 3. Myo1^S742^ phosphorylation fluctuates in a cell cycle dependent manner.** A *cdc10.v50* culture was synchronized in G1 by shifting to 36°C for 240 min before returning to 25°C at time 0. Samples of cells were taken every 20 minutes from the release and processed for western blotting to monitor of Myo1^S742^ phosphorylation (A). The membrane was subsequently probed with anti-Myo1 antibodies (B) to monitor total Myo1. Equal loading was monitored by Ponceau staining of the membrane. (C) Densitometry measurements of the bands in these blots are plotted along with the % of cells in the culture with septa.

**Supplementary Figure 4. Cam1 and Cam2 do not interact directly.** (A) Overlaid OD280 spectra were recorded from eluate from a Superdex 75 gel filtration column which had been loaded with either Cam1 (grey line), Cam2 (black line) or Cam1 and Cam2 (red line) under identical 4 mM EGTA buffer conditions. (B) Maximum IAANS fluorescence values (440 nm) of 0.5 μM Cam1-IAANS at a range of pCa values. Black symbols show values of Cam1-IAANS, red symbols show values of Cam1-IAANS with 5 μM Cam2 protein. 2 mM Ca-EGTA buffers were used to give indicated pCa values. pCa50 values calculated from Origin fitting analysis - Hill equation.

**Supplementary Figure 5. Multiple labelling strategies for Myo1, Cam1 and Cam2 disrupts normal distribution.** Cam1 has increased cytoplasmic signal and reduced signal at endocytic foci in cells expressing both *cam1.gfp* and *mCherry.myo1* (A (GFP-green, mCherry-magenta)) compared to cells expressing *cam1.gfp* alone (B). Similarly, Cam1 has increased cytoplasmic signal and reduced relative signal at endocytic foci in *CFP-myo1 cam1.mCherry* cells (C). (D) Growth curves of prototroph *cam1.gfp* (green) and *cam1.gfp mCherry.myo1* cells cultured in EMMG at 25 °C. (E) Cam1 (green) localisation is disrupted in *cam1.gfp cam2.mCherry* cells, with less Cam1 on endocytic foci, and localising to the mitotic spindle which is never observed in cells expressing FP labelled Cam1 alone.

**Supplementary Movie 1:** Timelapse of TIRFM imaged *mNeongreen.myo1* cells showing rapid single molecule interactions of Myo1 at the plasma membrane. Frame Rate: 15 msec / frame.

**Supplementary Movie 2:** Timelapse of TIRFM imaged *mNeongreen.myo1* cells showing endocytosis associated interactions of Myo1 at the plasma membrane. Frame rate: 50 msec / frame.

**Supplementary Movie 3:** Timelapse of TIRFM imaged *cam2.gfp* cells showing Cam2 recruiting to endocytic vesicles, to which it remains associated after scission and internalisation of the endosome. Frame rate: 50 msec / frame.

**Supplementary Movie 4:** Timelapse of TIRFM imaged *cam1.mCherry cam2.gfp* cells showing early recruitment of Cam1 (red) subsequent recruitment of Cam2 (green) to sites of endocytosis. Cam1 disassociates prior to vesicle scission, while Cam2 remains associated with the internalised endosome. Frame rate: 50 msec / frame.

**Supplementary Movie 5:** Timelapse of maximum projections from 13-z slice widefield images of *mNeongreen.myo1* cells showing typical examples of Myo1 dynamics in vegetative and meiotic cells. Frame rate: 650 msec / frame.

**Supplementary Movie 6:** Timelapse of maximum projections from 13-z slice widefield images of *cam1.gfp* cells showing typical examples of Cam1 dynamics in vegetative and meiotic cells. Frame rate: 650 msec / frame.

**Supplementary Movie 7:** Timelapse of maximum projections from 13-z slice widefield images of *cam2.gfp* cells showing typical examples of Cam2 dynamics in vegetative and meiotic cells. Frame rate: 650 msec / frame.

**Supplementary Movie 8:** Timelapse of maximum projections from 13-z slice widefield images of *gfp.act1* cells showing typical examples of Act1 dynamics in vegetative and meiotic cells. Frame rate: 650 msec / frame.

**Supplementary Table 1:** Strains used during this study.

**Supplementary Table 2:** Oligonucleotides used during this study.

