## Supplementary material for "TORC2 dependent phosphorylation modulates calcium regulation of fission yeast myosin"

Figure S1:

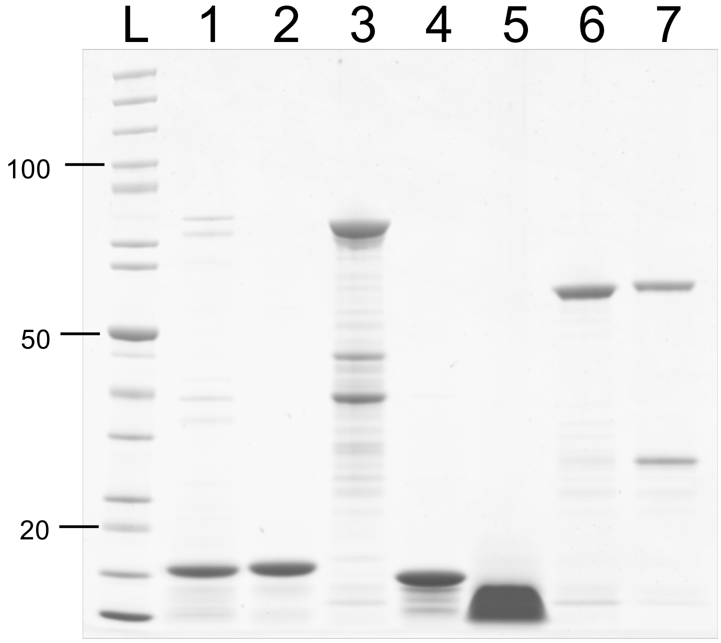


L – Ladder
1 – Cam1-ACE
2 – Cam1-IAANS
3 – Cam1-FRET
4 – Cam2
5 – IQ12 peptide – not used in study
6 – Myo1IQ12 FRET
7 – Myo1IQ12-S742D FRET

Figure S2:

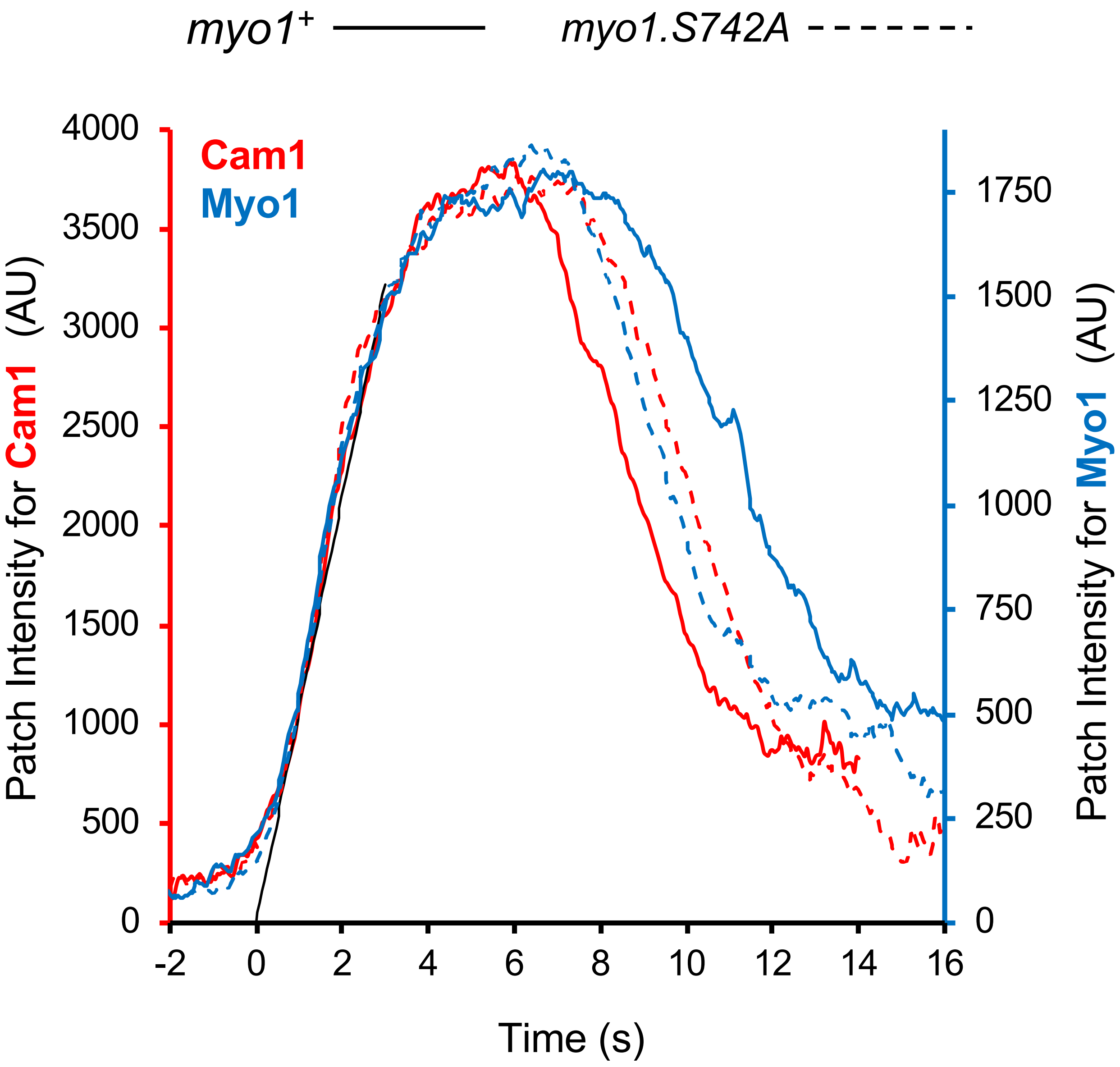

*myo1*+
*myo1.S742A*
Cam1
Myo1
Patch Intensity for Cam1 (AU)
Patch Intensity for Myo1 (AU)
Time (s)
4000
3500
3000
2500
2000
1500
1000
500
0
1750
1500
1250
1000
750
500
250
0
-2
0
2
4
6
8
10
12
14
16

Figure S3:

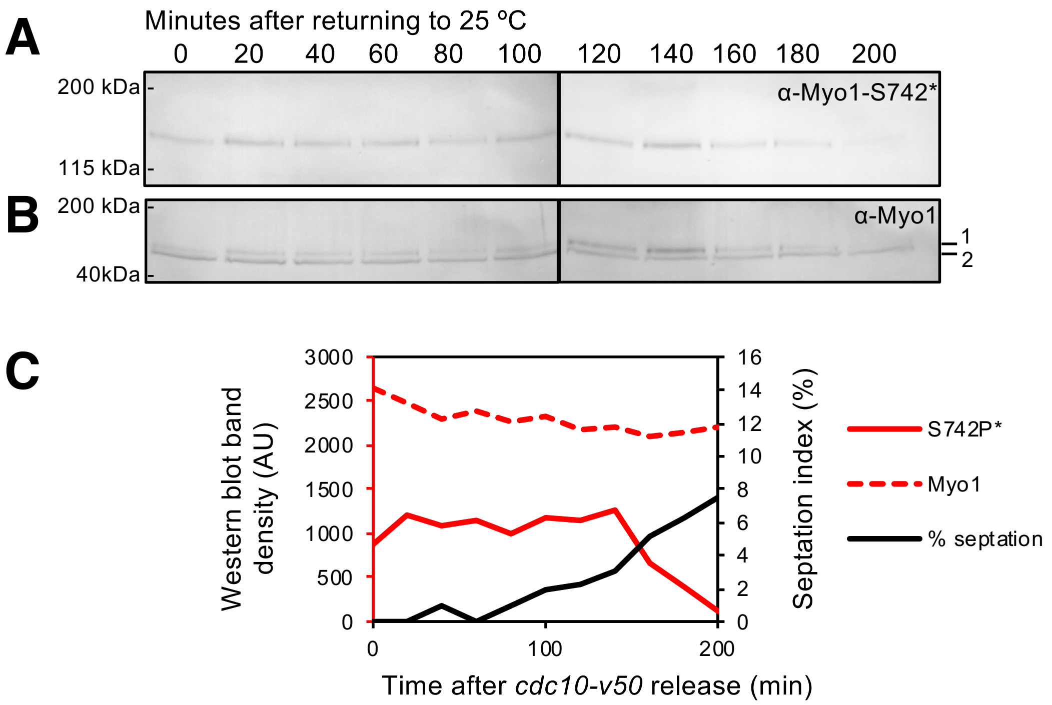
A
Minutes after returning to 25 °C
0 20 40 60 80 100 120 140 160 180 200
200 kDa
115 kDa
α-Myo1-S742*
B
200 kDa
40kDa
α-Myo1
1
2
C
Western blot band density (AU)
3000
2500
2000
1500
1000
500
0
Septation index (%)
16
14
12
10
8
6
4
2
0
0
100
200
Time after *cdc10-v50* release (min)
S742P*
Myo1
% septation

Figure S4:

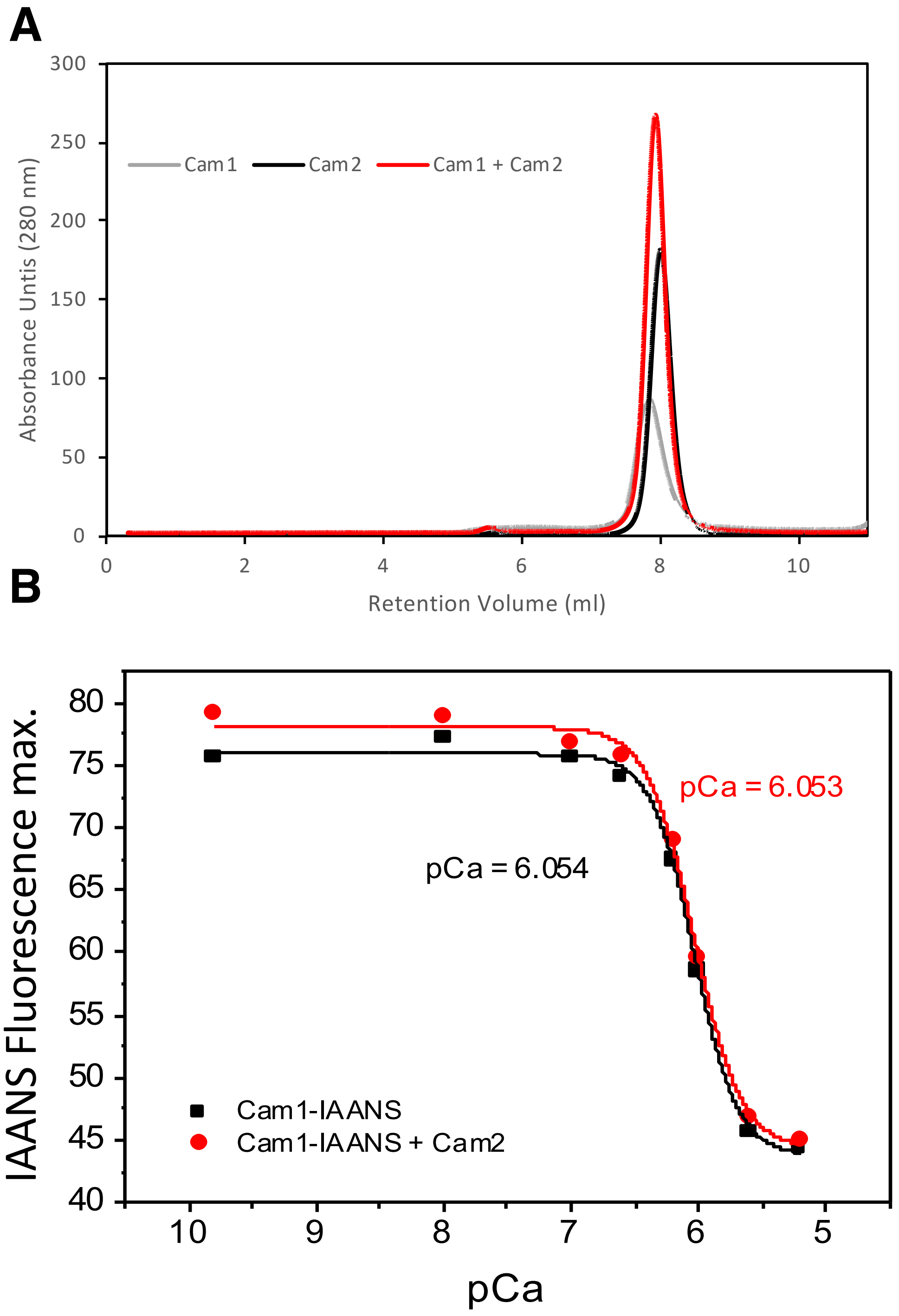
A
B
Cam1
Cam2
Cam1 + Cam2
Absorbance Untis (280 nm)
Retention Volume (ml)
pCa = 6.053
pCa = 6.054
Cam1-IAANS
Cam1-IAANS + Cam2
IAANS Fluorescence max.
pCa

Figure S5:

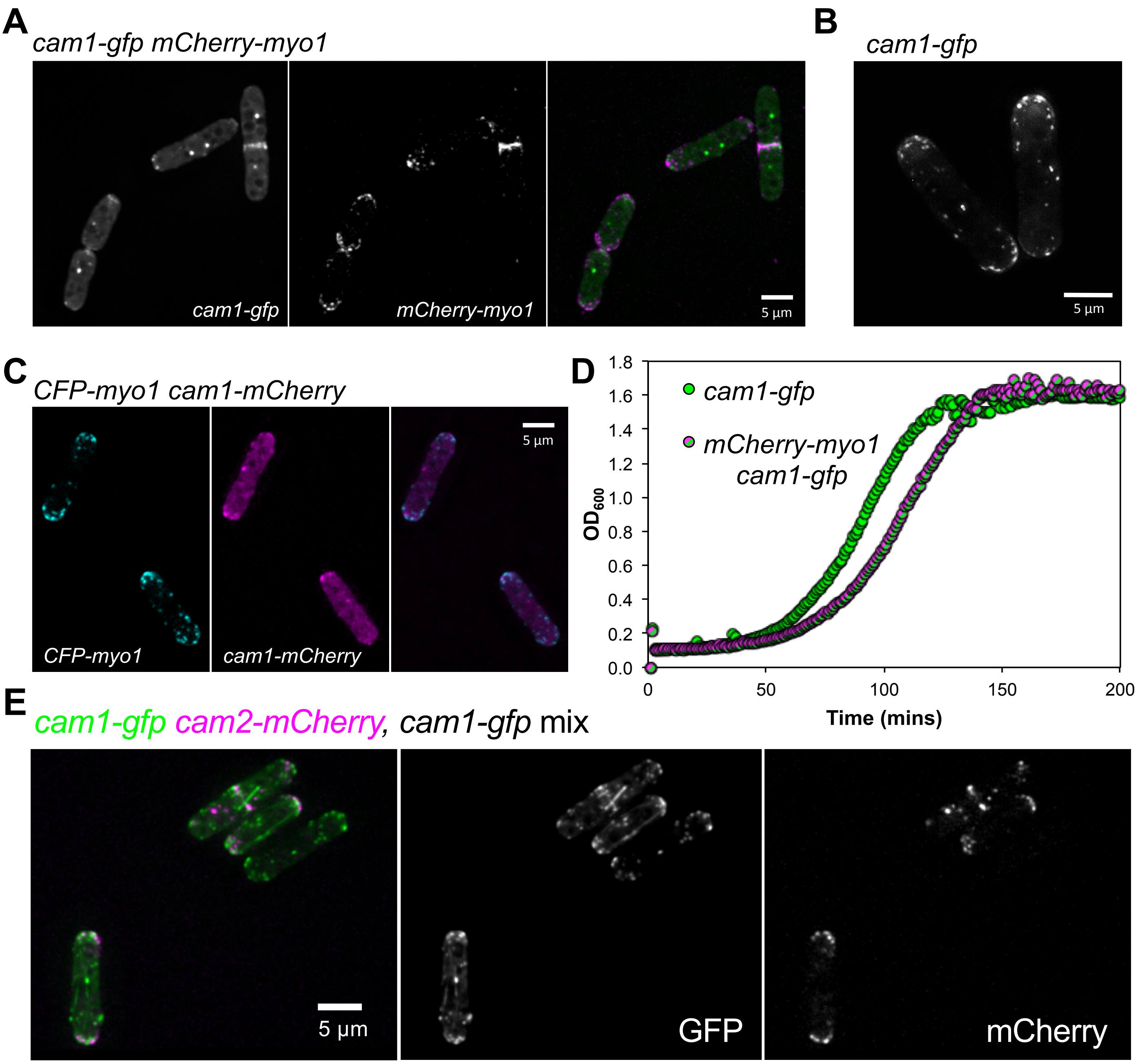
A
cam1-gfp mCherry-myo1
cam1-gfp
mCherry-myo1
5 µm
B
cam1-gfp
5 µm
C
CFP-myo1 cam1-mCherry
CFP-myo1
cam1-mCherry
5 µm
D
cam1-gfp
mCherry-myo1 cam1-gfp
OD600
Time (mins)
0.0
0.2
0.4
0.6
0.8
1.0
1.2
1.4
1.6
1.8
0
50
100
150
200
E
cam1-gfp cam2-mCherry, cam1-gfp mix
5 µm
GFP
mCherry
