## Supplementary material for "TORC2 dependent phosphorylation modulates calcium regulation of fission yeast myosin"

**Supplementary Table 1**: *Strains used during this study.*

| Strain | Source |
|---|---|
| wild type $h^-$ | Lab stock |
| *cdc10.v50* $h^-$ | (Nurse et al, 1976) |
| *cdc25.22* $h^-$ | (Nurse et al, 1976) |
| *myo1::kanMX6* | (Sirotkin *et al*, 2005) |
| *cam2::URA4 ura4.d18* $h^-$ | (Itadani *et al*, 2007) |
| *sla2.mCherry:natMX6* $h^-$ | (Alvarez-Tabarés *et al*, 2007) |
| *leu1::nmt81gfp.act1:URA4 ura4.d18* $h^{90}$ | (Doyle *et al*, 2009) |
| *LifeACT.mCherry:LEU2 leu1.32* $h^-$ | (Huang *et al*, 2012) |
| *acp1.gfp:kanMX6* $h^-$ | (Baker *et al*, 2016) |
| *ste20::kanMX6* $h^-$ | (Baker et al, 2016) |
| *gad8::kanMX6* $h^-$ | (Baker et al, 2016) |
| *myo1.S742A:URA4 ura4.d18* $h^-$ | This study |
| *myo1.S742D:URA4 ura4.d18* $h^-$ | This study |
| *mNeongreen.myo1:URA4 ura4.d18* $h^-$ | This study |
| *mNeongreen.myo1.S742A:URA4 ura4.d18* $h^-$ | This study |
| *mNeongreen.myo1:URA4 ura4.d18* $h^{90}$ | This study |
| *mNeongreen.myo1.S742A:URA4 ura4.d18* $h^{90}$ | This study |
| *mNeongreen.myo1:URA4 cam2::URA4 ura4.d18* $h^-$ | This study |
| *yfp.myo1:kanMX6 sid4.tdTomato:hphMX6* | This study |
| *cam1.gfp:kanMX6* $h^-$ | This study |
| *cam1.gfp:kanMX6 myo1::kanMX6* $h^-$ | This study |
| *cam1.gfp:kanMX6 myo1.S742A:URA4 ura4.d18* $h^-$ | This study |
| *cam1.gfp:kanMX6* $h^{90}$ | This study |
| *cam1.gfp:kanMX6 myo1.S742A:URA4 ura4.d18* $h^{90}$ | This study |
| *cam2.gfp:kanMX6* $h^-$ | This study |
| *cam2.gfp:kanMX6 myo1::kanMX6* $h^-$ | This study |
| *cam2.gfp:kanMX6 myo1.S742A:URA4 ura4.d18* $h^-$ | This study |
| *cam1.gfp:kanMX6 cam2::URA4 ura4.d18* $h^-$ | This study |
| *cam2.gfp:kanMX6* $h^{90}$ | This study |
| *cam2.gfp:kanMX6 myo1.S742A:URA4 ura4.d18* $h^{90}$ | This study |
| *leu1::nmt41gfp.myo1:URA4 ura4.d18 cam1.mCherry:hphMX6* $h^-$ | This study |
| *cam1.mCherry:hphMX6 cam2.gfp:kanMX6* $h^-$ | This study |
| *sla2.mCherry:natMX6 myo1.S742A:URA4 ura4.d18* $h^-$ | This study |
| *LifeACT.mCherry:LEU2 leu1.32 myo1::kanMX6* $h^-$ | This study |
| *LifeACT.mCherry:LEU2 leu1.32* $h^{90}$ | This study |
| *myo1.S742A:URA4 ura4.d18 sla2.mCherry:natMX6* $h^-$ | This study |

**Supplementary Table 2**: *Oligonucleotides used during this study.*

| Olig # | Name | Sequence (5'-3') |
|---|---|---|
| 226 | 5'Nde1cam1 | CATATGACTACCCGTAACC |
| 227 | 3'BamH1cam1 | GGATCCCTACTTGGAAGAAATG |
| 393 | 5'Nde1cam2 | CATATGCCTGCCTCCAAAGAACAAACCG |
| 394 | 3'BamH1cam2 | GGATCCCTATTTTGCCATGATTCTCTG |
| 403 | 5'Xho1YPet | CTCGAGATGGTGAGCAAAGGCGAAGAGC |
| 404 | 3'BamH1H6G3YPet | GGATCCTTAATGATGATGATGATGATGACCACCACCCTTATAG<br>AGCTCGTTCATGCCC |
| 405 | 5'Nde1CyPet | CATATGGTGAGCAAGGGAGAGG |
| 406 | 3'CyPetMyo1IQ1Xho1 | CTCGAGAGCTTCAGATCTTCTTCTAACATAAGAACGCCAAGCA<br>CGTTGTATACGGGTTGCCATGTCGACTTTGTACAGTTCGTCCA<br>TGCCGTGGGTG |
| 425 | BglII Myo1IQ12 Xho1F | GATCCGAAGCTGCTGCTTGTATTCAGAAGTTGTGGAATAGGAA<br>CAAAGTTAACATGGAACTTGAAC |
| 426 | BglII Myo1IQ12 Xho1R | TCGAGTTCAAGTTCCATGTTAACTTTGTTCCTATTCCACAACTT<br>CTGAATACAAGCAGCAGCTTCG |
| 427 | IQ12 S742D F | CTTATGTTAGACGTCGCGACGAAGCTGCTGCTTG |
| 428 | IQ12 S742D R | CAAGCAGCAGCTTCGTCGCGACGTCTAACATAAG |
| 429 | Sal1Myo1IQ2BglIIF | TCGACGCTGCTGCTTGTATTCAGAAGTTGTGGAATAGGAACAA<br>AGTTAACATGGAACTTGAACTCGAGA |
| 430 | Sal1Myo1IQ2BglIIR | GATCTCTCGAGTTCAAGTTCCATGTTAACTTTGTTCCTATTCCA<br>CAACTTCTGAATACAAGCAGCAGCG |

Note: Underscored sequences denotes endonuclease recognition sites
